# Recombination and mutational robustness in neutral fitness landscapes

**DOI:** 10.1101/556134

**Authors:** Alexander Klug, Su-Chan Park, Joachim Krug

## Abstract

Mutational robustness quantifies the effect of random mutations on fitness. When mutational robustness is high, most mutations do not change fitness or have only a minor effect on it. From the point of view of fitness landscapes, robust genotypes form neutral networks of almost equal fitness. Using deterministic population models it has been shown that selection favors genotypes inside such networks, which results in increased mutational robustness. Here we demonstrate that this effect is massively enhanced by recombination. Our results are based on a detailed analysis of mesa-shaped fitness landscapes, where we derive precise expressions for the dependence of the robustness on the landscape parameters for recombining and non-recombining populations. In addition, we carry out numerical simulations on different types of random holey landscapes as well as on an empirical fitness landscape. We show that the mutational robustness of a genotype generally correlates with its recombination weight, a new measure that quantifies the likelihood for the genotype to arise from recombination. We argue that the favorable effect of recombination on mutational robustness is a highly universal feature that may have played an important role in the emergence and maintenance of mechanisms of genetic exchange.

**Author summary:** Two long-standing and seemingly unrelated puzzles in evolutionary biology concern the ubiquity of sexual reproduction and the robustness of organisms against genetic perturbations. Using a theoretical approach based on the concept of a fitness landscape, in this article we argue that the two phenomena may in fact be closely related. In our setting the hereditary information of an organism is encoded in its genotype, which determines it to be either viable or non-viable, and robustness is defined as the fraction of mutations that maintain viability. Previous work has demonstrated that the purging of non-viable genotypes from the population by natural selection leads to a moderate increase in robustness. Here we show that genetic recombination acting in combination with selection massively enhances this effect, an observation that is largely independent of how genotypes are connected by mutations. This suggests that the increase of robustness may be a major driver underlying the evolution of sexual recombination and other forms of genetic exchange throughout the living world.

## Introduction

The reshuffling of genetic material by recombination is a ubiquitous part of the evolutionary process across the entire range of organismal complexity. Starting with viruses as the simplest evolving entities, recombination occurs largely at random during the coinfection of a cell by more than one virus strain [1]. For bacteria the mechanisms involved in recombination are already more elaborate and present themselves in the form of transformation, transduction and conjugation [2, 3]. In eukaryotic organisms, sexual reproduction is a nearly universal feature, and recombination is often a necessary condition for the creation of offspring. Although its prevalence in nature is undeniable, the evolution and maintenance of sex is surprising since compared to an asexual population, only half of a sexual population is able to bear offspring and additionally a suitable partner needs to be found [4, 5]. Whereas the resulting two-fold cost of sex applies only to organisms with differentiated sexes [6], the fact that genetic reshuffling may break up favorable genetic combinations or introduce harmful variants into the genome poses a problem also to recombining microbes that reproduce asexually [7, 8]. Since this dilemma was noticed early on in the development of evolutionary theory, many attempts have been undertaken to identify evolutionary benefits of sex and recombination based on general population genetic principles [9–17].

In this article we approach the evolutionary role of recombination from the perspective of fitness landscapes. The fitness landscape is a mapping from genotype to fitness, which encodes the epistatic interactions between mutations and provides a succinct representation of the possible evolutionary trajectories [18]. Previous computational studies addressing the effect of recombination on populations evolving in epistatic fitness landscapes have revealed a rather complex picture, where evolutionary adaptation can be impeded or facilitated depending on, e.g., the structure of the landscape, the rate of recombination or the time frame of observation [19–24].

Here we focus specifically on the possible benefit of recombination that derives from its ability to enhance the mutational robustness of the population. A living system is said to be robust if it is able to maintain its function in the presence of perturbations [25–29]. In the case of mutational robustness these perturbations are genetic, and the robustness of a genotype is quantified by the number of mutations that it can tolerate without an appreciable change in fitness. Robust genotypes that are connected by mutations therefore form plateaux in the fitness landscape that are commonly referred to as neutral networks [30–32]. Mutational robustness is known to be abundant at various levels of biological organization, but its origins are not well understood. In particular, it is not clear if mutational robustness should be viewed as an evolutionary adaptation, or rather reflects the intrinsic structural constraints of living systems.

An argument in favor of an adaptive origin of robustness was presented by van Nimwegen and collaborators, who showed that selection tends to concentrate populations in regions of a neutral network where robustness is higher than average [30]. Whereas this result is widely appreciated, the role of recombination for the evolution of robustness has received much less attention. An early contribution that can be mentioned in this context is due to Boerlijst *et al.* [33], who discuss the error threshold in a viral quasi-species model with recombination and point out in a side note that *“in sequence space recombination is always inwards pointing.”* This observation was picked up by Wilke and Adami [34] in a review on the evolution of mutational robustness, where they conjecture that the enhancement of robustness by selection should be further amplified by recombination, because “*recombination alone always creates sequences that are within the boundaries of the current mutant cloud*.” In fact a clear indication of this effect had been reported somewhat earlier by Xia and Levitt in a computational study of the evolution of 2D lattice proteins in the presence and absence of recombination [35]. The native folding structure of a given sequence is determined by its global minimum free energy. Due to the restricted number of attainable folds, most structures are degenerate in the sense that many sequences fold into the same structure. These sequences form neutral networks in sequence space. Xia and Levitt [35] consider two scenarios, in which evolution is dominated by mutation and by recombination, respectively. The results show that in the latter case the concentration of thermodynamically stable protein sequences is enhanced, which is qualitatively explained by the fact that recombination tends to focus the sequences near the center of their respective neutral network. Therefore most often a single mutation does not change the folding structure.

More recently, Azevedo *et. al* [36] used a model of gene regulatory networks to investigate the origin of negative epistasis, which is a requirement for the advantage of recombination according to the mutational deterministic hypothesis [13]. In this study a gene network is encoded by a matrix of interaction coefficients. It is defined to be viable if its dynamics converges to a stable expression pattern and non-viable otherwise. Thus the underlying fitness landscape is again neutral. Based on their simulation results the authors argue that recombination of interaction matrices reduces the recombinational load, which in turn leads to an increase of mutational robustness and induces negative epistasis as a byproduct. In effect, then, recombination selects for conditions that favor its own maintenance. Other studies along similar lines have been reviewed in [37]. Taken together they suggest that the positive effect of recombination on robustness may be largely independent of the precise structure of the space of genotypes or the genotype-phenotype map. Indeed, a related scenario has also been described in the context of computational evolution of linear genetic programs [38].

Finally, in a numerical study that is similar to ours in spirit, Szöllősi and Derényi considered the effect of recombination on the mutational robustness of populations evolving on different types of neutral fitness landscapes [39]. Using neutral networks that were either generated at random or based on RNA secondary structure, they found that recombination generally enhances mutational robustness by a significant amount. Moreover, they showed that this observation holds not only for infinite populations but also for finite populations, as long as these are sufficiently polymorphic. The goal of this article is to explain these scattered observations in a systematic and quantitative way. For this purpose we begin by a detailed examination of the simplest conceivable setting consisting of a haploid two-locus model with three viable and one lethal genotype [32]. We derive explicit expressions for the robustness as a function of the rates of mutation and recombination that demonstrate the basic phenomenon and guide the exploration of more complex situations. The two-locus results are then generalized to mesa landscapes with *L* diallelic loci, where genotypes carrying up to *k* mutations are viable and of equal fitness [40–43]. Subsequently two types of random holey landscape models are considered, including a novel class of sea-cliff landscapes in which the fraction of viable genotypes depends on the distance to a reference sequence. A unified picture of the effect of recombination in neutral landscapes is developed through the concept of the recombination weight, which is a measure for the likelihood of a genotype to arise from a recombination event and generally correlates with mutational robustness. Using an empirical fitness landscape as an example, we demonstrate how the recombination weight allows one to quantify the competition between selection and recombination as a function of recombination rate. Throughout we describe the evolutionary dynamics by a deterministic, discrete time model that will be introduced in the next section.

## Models and Methods

### Genotype space

We consider a haploid genome with *L* loci and the corresponding genotype is represented by a sequence *σ* = (*σ*_1_, *σ*_2_,*…, σ*_*L*_) of length *L*. The index *i* labels genetic loci and each locus carries an allele specified by *σ*_*i*_. Here we rely on binary sequences, which means that there are only two different alleles *σ*_*i*_ ∈ {0, 1}. This can be either seen as a simplification in the sense that only two alleles are assumed to exist, or in the sense that the genome consisting of all zeros describes the wild type, and the 1’s in the sequence display mutations for which no further distinctions are made.

The resulting genotype space is a hypercube of dimension *L*, where the 2^*L*^ genotypes represent vertices, and two genotypes that differ at a single locus and are mutually reachable by a point mutation are connected by an edge. A metric is introduced by the Hamming distance

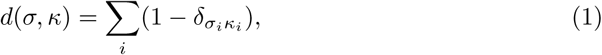

which measures the number of point mutations that separate two genotypes *σ* and *κ*. Here and in the following the Kronecker symbol is defined as *δ*_*xy*_ = 1 if *x* = *y* and *δ*_*xy*_ = 0 otherwise. The genotype 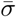 at maximal distance 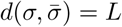 from a given genotype *σ* is called its antipodal, and can be defined by 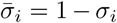. Finally, in order to generate a fitness landscape, a (Wrightian) fitness value *w*_*σ*_ is assigned to each genotype.

### Dynamics

The forces that drive evolution are selection, mutation and recombination. To model the dynamics we assume a discrete and non-overlapping generation model, of Wright-Fisher type. For simplicity, the population size is assumed to be infinite, which implies that demographic stochasticity, or genetic drift, is absent. Numerical simulations of evolution on neutral networks have shown that the infinite population dynamics is already observable for moderate population sizes, which justifies this approximation [30, 39, 44].

Once the frequency *f*_*σ*_(*t*) of a genotype *σ* at generation *t* is given, the frequency at the next generation is determined in three steps representing selection, mutation, and recombination. After the selection step, the frequency *q*_*σ*_(*t*) is given as

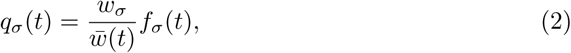

where 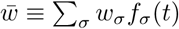 is the mean population fitness at generation *t*. After the mutation step, the frequency *p*_*σ*_(*t*) is given as

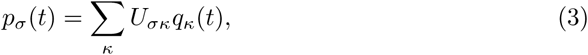

where *U*_*σκ*_ is the probability that an individual with genotype *κ* mutates to genotype *σ* in one generation. Here, we assume that alleles at each locus mutate independently, and the mutation probability *µ* is the same in both directions (0 → 1 and 1 → 0) and acrosloci. This leads to the symmetric mutation matrix

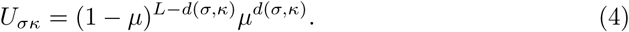

In order to incorporate recombination we have to consider the probability that two parents with respective genotypes *κ* and *τ* beget a progeny with genotype *σ* by recombination. This is represented by the following equation:

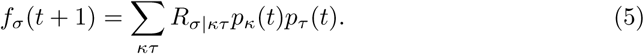

Descriptively speaking, two genotypes (*κ* and *τ)* are taken to recombine with a probability that is equal to their frequency in the population (after selection and mutation). The probability for the offspring genotype *σ* is then given by *R*_*σ*|*κτ*_. These probabilities depend of course on the parent genotypes *κ* and *τ* but also on the recombination scheme. Here we consider a uniform and a one-point crossover scheme; see Fig 1 for a graphical representation. These two represent extremes in a spectrum of possible recombination schemes. Nevertheless we will show that both lead to qualitatively similar results in the regimes of interest. In the case of uniform crossover the recombination probabilities are given by

**Fig 1.**
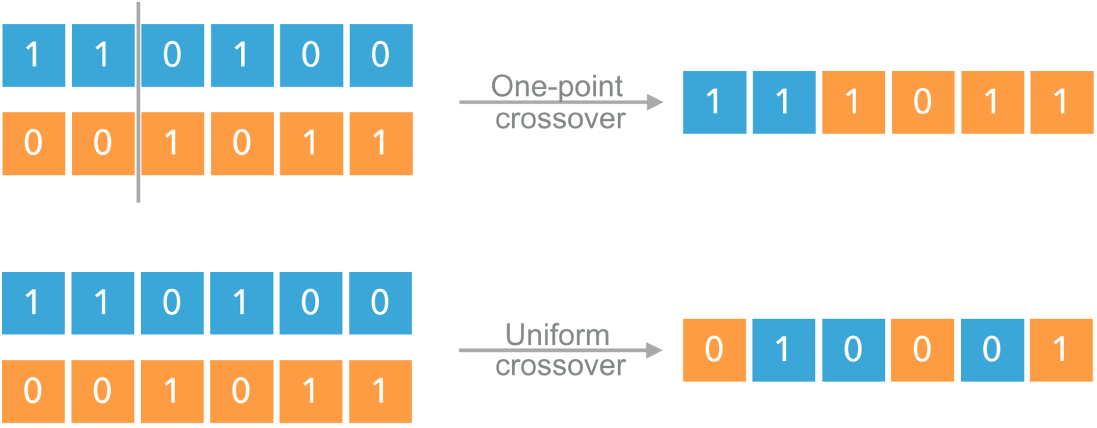
Recombination schemes. In the one-point crossover scheme, the parent genotypes are cut once between two randomly chosen loci and recombined to form the offspring. In the uniform crossover scheme, at each locus of the offspring, an allele present in one of the parents is chosen at random.

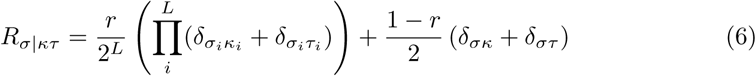

and in the case of one point crossover the probabilities can be written as

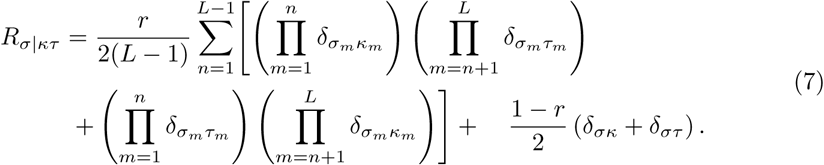

In both equations a variable *r* ∈ (0, 1) appears which describes the recombination rate. For *r* = 0 no recombination occurs and *f*_*σ*_(*t* + 1) is the same as *p*_*σ*_(*t*). For *r* = 1 recombination is a necessary condition for the creation of offspring (obligate recombination). But also intermediate values of *r* can be chosen as they occur in nature, e.g., for bacteria and viruses.

In the following we are mostly interested in the equilibrium frequency distribution 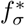 of a population, which is determined by the stationarity condition

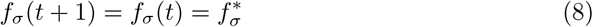

for all genotypes *σ*.

### Mutational robustness

From the point of view of fitness landscapes the occurrence of mutational robustness implies that fitness values of neighboring genotypes are degenerate, giving rise to neutral networks in genotype space [27, 30–32]. In order to model this situation we use two-level landscapes that only differentiate between genotypes that are viable (*w*_*σ*_ = 1) or lethal (*w*_*σ*_ = 0). Any selective advantage between viable genotypes is assumed to be negligible. The mutational robustness of a population can then be measured by the average fraction of viable point mutations in an individual, which depends on the population distribution in genotype space [30, 31]. It increases if the population mainly adapts to genotypes for which most point mutations are viable. Therefore we define mutational robustness *m* as the average fraction of viable point mutations of a population,

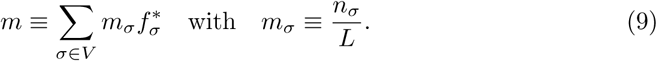

Here the sum is over the set *V* of all viable genotypes and *n*_*σ*_ is the number of viable point mutations of genotype *σ*. We will refer to *m*_*σ*_ as the mutational robustness of the genotype. The expression is normalized by the total number of loci *L*, since in an optimal setting the entire population has *L* viable genotypic neighbors and *m*_*σ*_ = 1 for all *σ* ∈ *V.* The value of *m* is thus constrained to be in the range [0, 1]. We weight the genotypes by their stationary frequencies 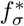, since we want to determine the mutational robustness of populations that are in equilibrium with their environment.

### Recombination weight

In order to elucidate the interplay of recombination and mutation robustness it will prove helpful to introduce a representation of how recombination can transfer genotypes into each other. The number of distinct genotypes that two recombining genotypes are able to create depends on their Hamming distance. In particular, the recombination of two identical genotypes does not create any novelty, whereas a genotype and its antipodal are able to generate all possible genotypes through uniform crossover.

Here we introduce a measure which expresses how many pairs of viable genotypes are able to recombine to a specific genotype. It is complementary to the mutational robustness, in the sense that instead of counting the viable mutation neighbors of a genotype, the size of its recombinational neighborhood of viable recombination pairs is determined. The recombinational neighborhood depends on the recombination scheme and the distribution of viable genotypes in the genotype space. For a given recombination scheme the probability for a genotype *σ* to be the outcome of recombination of two genotypes *κ, τ* is given by the recombination tensor *R*_*σ*|*κτ*_. The *recombination weight λ*_*σ*_ is therefore obtained by summing the recombination tensor over all ordered pairs of viable genotypes,

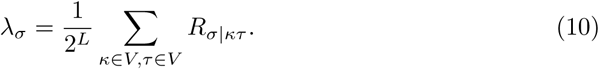

It can be seen from (5) that *λ*_*σ*_ = 1 when all genotypes are viable, and hence the normalization by 2^*L*^ ensures that the recombination weight lies in the range [0, 1]. Under this normalization, the recombination weights sum to 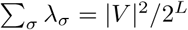, where |*V* | stands for the number of viable genotypes. In the following the genotype maximizing *λ*_*σ*_ will be referred to as the *recombination center* of the landscape.

Since neutral landscapes only differentiate between viable (unit fitness) and lethal (zero fitness) genotypes, the recombination weight (10) can alternatively be written as a sum over all ordered pairs of genotypes whereby the recombination tensor is multiplied by the pair’s respective fitness,

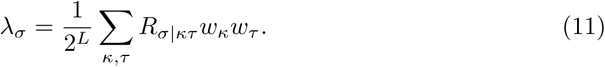

In this way the concept naturally generalizes to arbitrary fitness landscapes. In the absence of recombination (*r* = 0) the recombination weight (11) of a genotype is simply proportional to its fitness, 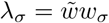, where 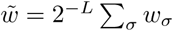 is the unweighted average fitness. Within our recombination schemes, the recombination tensor depends linearly on *r* and, by definition, so does the recombination weight. Accordingly, for general *r* the recombination weight interpolates linearly between the limiting values at *r* = 0 and *r* = 1. Since *λ*_*σ*_ for *r* = 0 is known, the remaining task will be to find *λ*_*σ*_ for *r* = 1.

### Visualization of fitness landscapes as networks

In order to visualize random neutral fitness landscapes with more than two loci we make use of a network representation, where genotypes that differ by a single mutation are connected by an edge. Nodes of the network then represent genotypes, which are arranged according to a spring layout that is based on a Fruchterman-Reingold force-directed algorithm [45]. To describe this algorithm briefly, nodes are made to repel each other, which is counteracted by edges that function as springs. This leads to a process of spring-force relaxation that arrives at an equilibrium state which in turn is used for the node positions. The equilibrium state is characterized by clustering of highly connected regions of nodes. Therefore this algorithm is only useful if not all nodes have the same number of edges. Hence edges attached to a lethal genotypes are deleted. This leads to a network in which only viable genotypes that differ by a single mutation are connected. Lethal genotypes are off the grid and create a ring of repelled nodes.

## Results

In the following sections we investigate how mutational robustness depends on the mutation and recombination rates. In order to test the generality of our results, we use, besides contrasting recombination schemes, also different neutral landscape models such as the mesa [40–43] and the percolation models [32, 46]. Additionally we introduce a more general landscape named sea-cliff model, which combines elements of both the landscape models and contains them as limiting cases. In the end, we discuss mutational robustness and its relation with recombination weight for an empirical landscape.

Two-locus models are commonly used in population genetics to gain a foothold in understanding evolutionary scenarios involving multiple recombining loci [32, 47–54]. Following this tradition, we first discuss a two-locus model and then extend our results to multi-locus models.

### Two-locus model

The simplest fitness landscape to study the mutational robustness of a population would be the haploid two-locus model in which all but one genotype are viable [32]; see Fig 2 for a graphical representation of the model. In this setting the population gains mutational robustness if the frequency of the genotype (0,0) for which both point mutations are viable increases relative to the genotypes (0,1) and (1,0). This model has been analyzed previously using a unidirectional mutation scheme where reversions 1 → 0 are suppressed [55, 56]. As a consequence, selection cannot contribute to mutational robustness because the genotype (0,0) goes extinct in the absence of recombination. Here we consider the case of bidirectional, symmetric mutations in which both selection and recombination contribute to robustness. A comparison of the two mutation schemes is provided in S1 Appendix.

**Fig 2.**
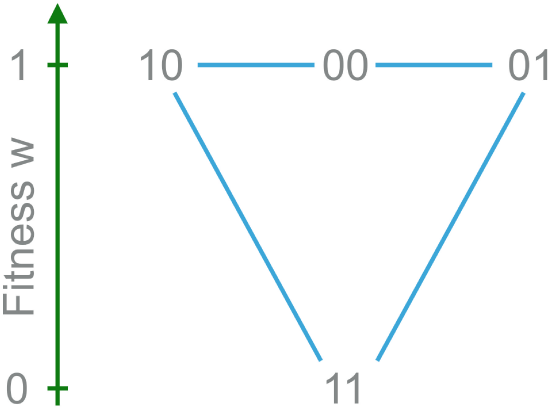
Two-locus model. Genotype (1,1) is lethal while the other three genotypes are viable with the same fitness. Here, genotype (0,0) is most robust since both its single mutants are viable.

We proceed to solve the equilibrium condition Eq (8). Since the equilibrium genotype frequencies 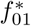 and 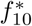 are the same due to the symmetry of the landscape and the mutation scheme, the recombination step at stationarity reads

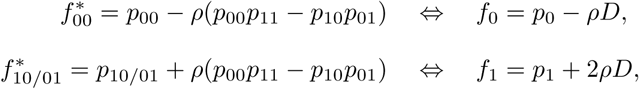

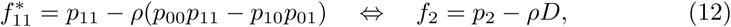

where *p*_*σ*_ is the (equilibrium) frequency of genotype *σ* after the mutation step, *f*_*i*_ and *p*_*i*_are the corresponding lumped frequencies [57] of all genotypes with *i* 1’s, and 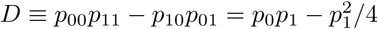 is the linkage disequilibrium after the mutation step. Notice that the one-point and uniform crossover schemes give the same equation form except that the parameter *ρ* is given by *ρ* = *r* in the case of one-point crossover and *ρ* = *r/*2 for uniform crossover. However, we would like to emphasize that this is a mere coincidence of the two-locus model which disappears as soon as *L* is larger than 2.

The lumped frequencies *q*_*i*_ of all genotypes with *i* 1’s after the selection step are given by

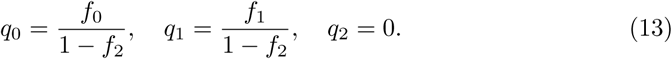

Applying the mutation step we obtain

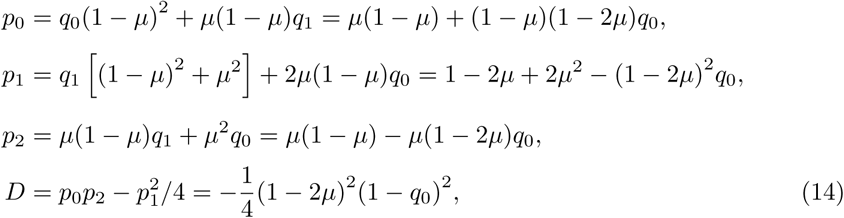

where we have used the normalization *q*_0_ + *q*_1_ = 1 to express the right hand sides in terms of *q*_0_. Putting everything together, the problem is reduced to solving the following third order polynomial equation for *q*_0_,

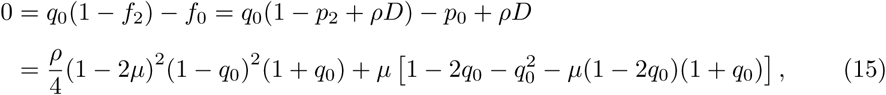

from which we can in principle find exact analytic expressions for 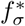. However, it is difficult to extract useful information from the exact solution. In the following we will therefore provide approximate solutions.

If we neglect recombination (*ρ* = 0), we obtain the following equilibrium genotype frequency distribution:

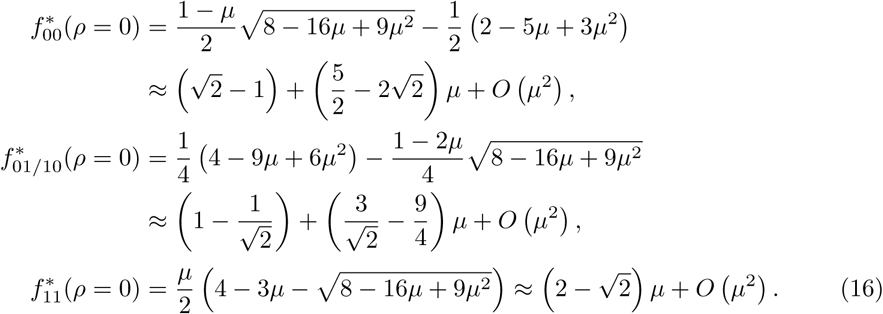

When *ρ* = 1, which corresponds to the one-point crossover scheme with *r* = 1, linkage equilibrium is restored after one generation [52]. Accordingly, we can treat each locus independently and get rather simple expressions for 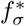 as

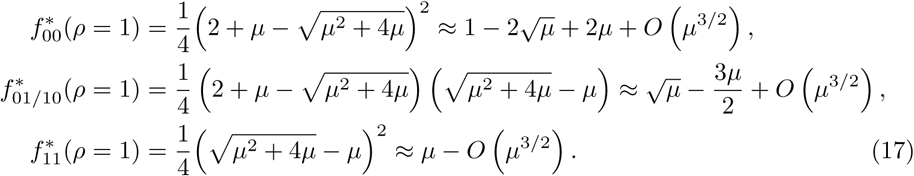

We depict the equilibrium solutions for the above two cases in Fig 3.

**Fig 3.**
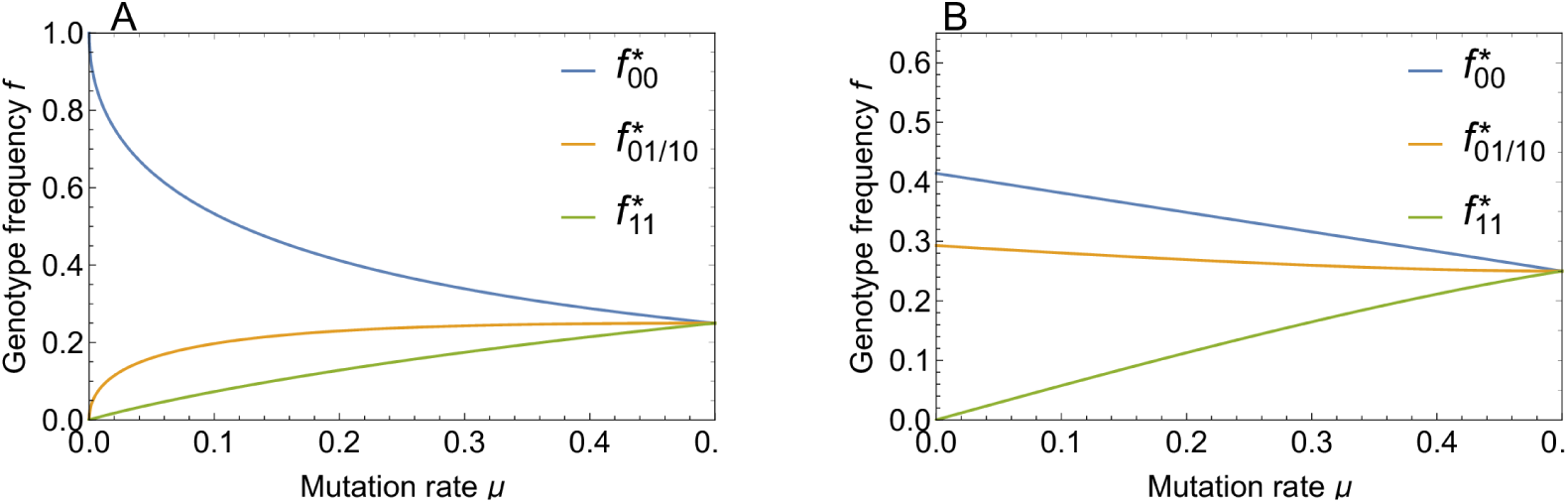
Equilibrium genotype frequencies in the two locus model. Genotype frequencies in the stationary state are shown as a function of mutation rate for (A) strong recombination (*ρ* = 1) and (B) no recombination (*ρ* = 0).

Now, the mutational robustness

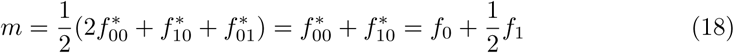

for the above two cases is obtained as

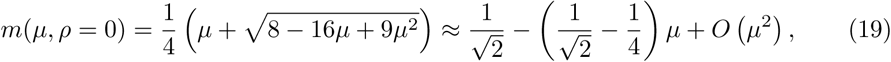

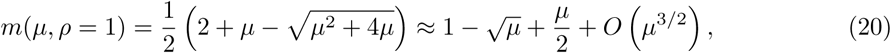

which is depicted in Fig 4. These results encapsulate in a simple form the main topic of this paper. Selection alone (*ρ* = 0) leads to a moderate increase of robustness from the baseline value 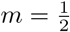 corresponding to a random distribution over genotypes, which is attained at 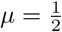, to 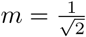 for *µ* → 0. In contrast, for recombining populations (*ρ* = 1) robustness is massively enhanced at small mutation rates due to the strong frequency increase of the most robust genotype (0,0) and reaches the maximal value *m* = 1 at *µ* = 0. The underlying mechanism is analogous to Kondrashov’s deterministic mutation hypothesis, which posits that recombination increases the genotypic variability and therefore makes selection more effective [13]. In the present case recombination increases the frequency of the double mutant genotype (1, 1), which is subsequently purged by selection, and thereby effectively drives the frequency of the allele 1 at both loci to zero. The enhancement of the frequency of the genotype (0,0) by recombination is also reflected in the recombination weights, which take on the values

**Fig 4.**
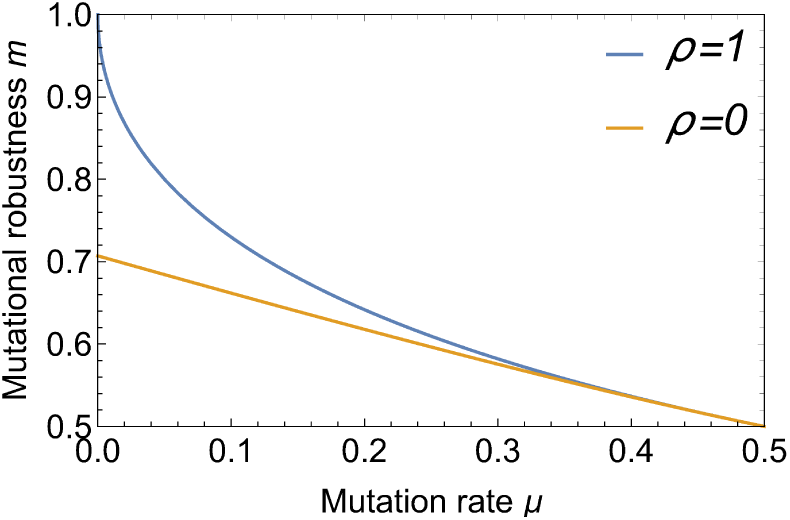
Mutational robustness as a function of mutation rate. The figure shows the robustness in the two-locus model at *ρ* = 0 and *ρ* = 1. Recombination leads to a massive enhancement of robustness for small mutation rates.

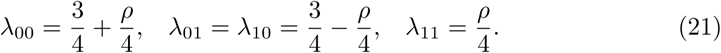

Thus the genotype (0,0) is the recombination center of the two-locus landscape.

Next we investigate how mutational robustness varies with *µ* for intermediate recombination rates, assuming that *µ* is small. As can be seen from Eq (15), the asymptotic behavior of the solution for small *ρ* and *µ* depends on which of the two parameters is smaller. We first consider the case *ρ* ≪ *µ* ≪ 1. Defining *l* = *ρ/*(4*µ*) ≪ 1, Eq (15) is approximated by

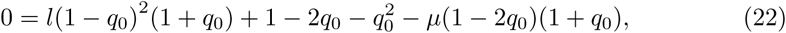

where we kept terms up to *O*(*µ*), since we have not determined whether *l* is smaller than *µ* or not. Since 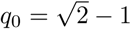 is the solution of Eq (22) for *l* = *µ* = 0, we set 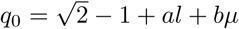 and solve the equation to leading order, which gives

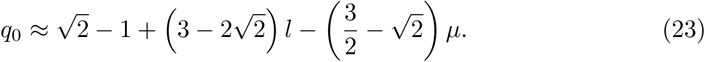

The mutational robustness then follows as

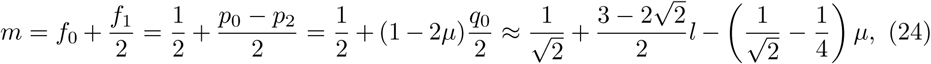

which is consistent with our previous result for *ρ* = 0; see Eq (19). We note that in this regime it is sufficient for the recombination rate to be of order *O*(*µ*^2^) to compensate the negative effect of mutations on mutational robustness, as the two effects cancel when *ρ* = *ρ*_*c*_ with

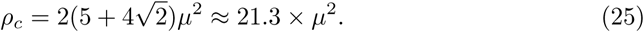

In the regime *ρ* ≫ *µ*, Eq (15) is approximated as

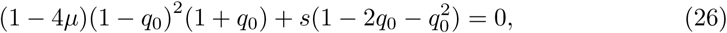

with *s* = 4*µ/ρ*. Again we have kept terms up to *O*(*µ*) because *µ* and *s* are of the same order if *ρ* = *O*(1). Since the solution of Eq (26) for *µ* = *s* = 0 is *q*_0_ = 1, we set *q*_0_ = 1 *− α* with *α* ≪ 1. Inserting this into Eq (26), we get 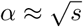. Since *α* ≫ *µ*,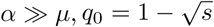 is the approximate solution to leading order. Hence

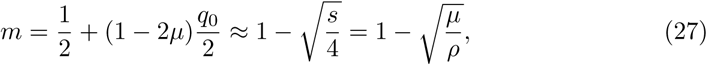

which is again consistent with our previous result for *ρ* = 1 in Eq (20). The square root dependence on *µ/ρ* derives from the corresponding behavior of the genotype frequency 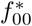 and has been noticed previously in the model with unidirectional mutations [55, 56].

For arbitrary *ρ* and *µ*, we have to use the full Eq (15). Fig 5 illustrates the behaviour of mutational robustness as a function of the recombination rate for different mutation rates and both recombination schemes. For small *µ*, a low rate of recombination suffices to bring the robustness close to its maximal value *m* = 1. More precisely, according to Eq (27), a robustness *m >* 1 *− ϵ* is reached for recombination rates *ρ > µ/c*^2^.

**Fig 5.**
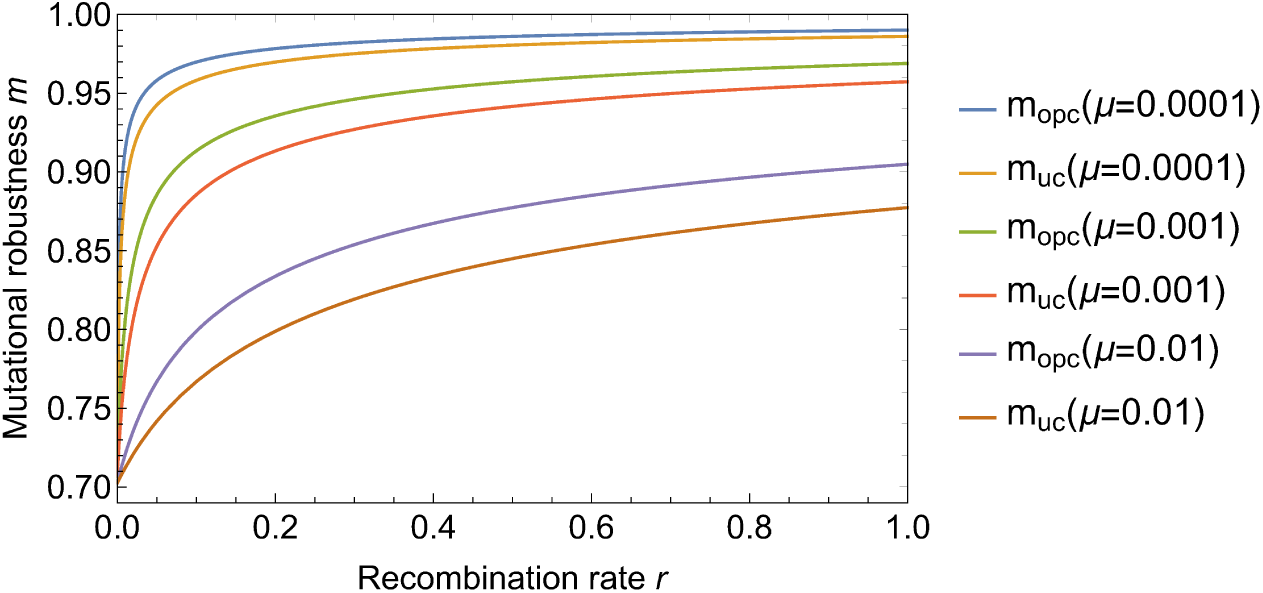
Mutational robustness as a function of recombination rate. The figure shows the mutational robustness for one-point crossover (*m*_opc_) and uniform crossover (*m*_uc_) and three different values of the mutation rate *µ*. When mutations are rare, a small amount of recombination is sufficient to significantly increase mutational robustness.

To summarize, we have seen that analytic results for the two-locus model are easily attainable. For multi-locus models it is much more challenging to derive analytical results, particularly in the presence of recombination. By way of contrast the dynamics induced only by mutation and selection are easier to understand: While mutations increase the genotype diversity in the population, fitter ones grow in frequency through selection, which reduces diversity. Although one might expect that recombination would increase diversity, a number of studies have shown that recombination is more likely to impede the divergence of populations. Recombining populations tend to cluster on single genotypes or in a limited region of a genotype space and furthermore the waiting times for peak shifts in multipeaked fitness landscapes diverge at a critical recombination rate [20, 24, 44, 51–53]. The results for the two-locus model presented above are consistent with this behaviour, as the genotype heterogeneity of the population decreases with increasing recombination rate (S1 Fig).

In the following we will investigate how the focusing effect of recombination enhances the mutational robustness of the population in three different multi-locus models.

### Mesa landscape

In the mesa landscape it is assumed that up to a certain number *k* of mutations all genotypes are functional and have unit fitness, whereas genotypes with more than *k* mutations are lethal and have fitness zero [43]. Hence the fitness landscape is defined as

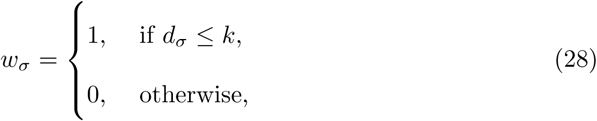

where *d*_*σ*_ is the Hamming distance to the wild-type sequence (0, 0,*…,* 0) or, equivalently, the number of loci with allele 1. We will refer to *k* as the mesa width or as the critical Hamming distance.

Such a scenario can for example be observed in the evolution of regulatory motifs, where the fitness depends on the binding affinity of the regulatory proteins and *d*_*σ*_ corresponds to the number of mismatches compared to the original binding motif [40, 42]. The two-locus model discussed in the preceding section corresponds to the mesa landscape with critical Hamming distance *k* = 1 and sequence length, *L* = 2. Here we ask to what extent the behavior observed for the two-locus model generalizes to longer sequences and variable *k*. Numerical simulations suggest that the strong increase of mutational robustness with recombination rate indeed persists in the general setting, and the particular recombination scheme seems to have only a minor influence; see Fig 6.

**Fig 6.**
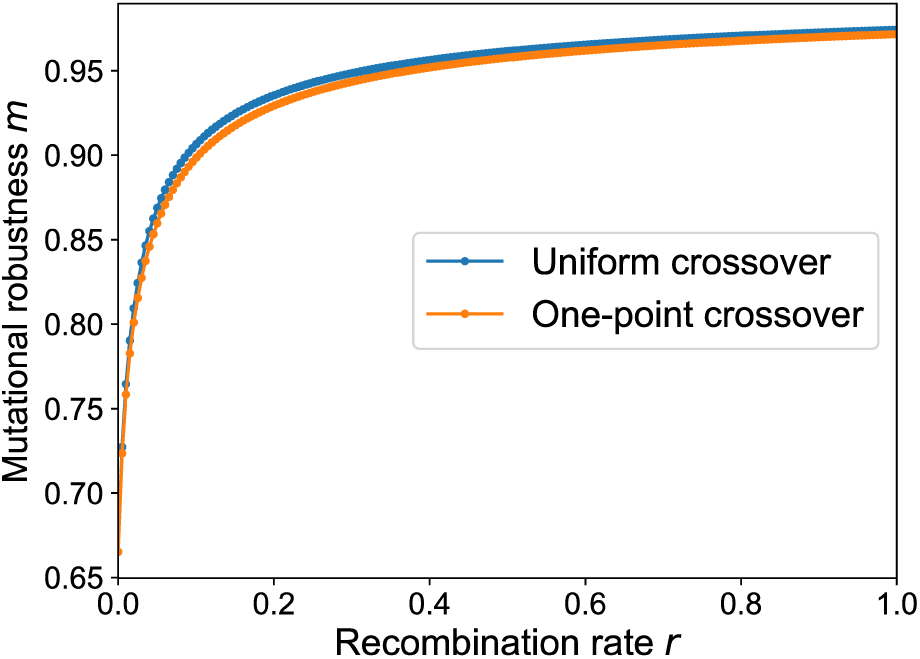
Mutational robustness in a mesa landscape as a function of recombination rate. Data points are obtained by numerically iterating the selection-mutation-recombination dynamics until the equilibrium state is reached. The parameters of the mesa landscape are *L* = 6, *k* = 2 and the mutation rate is *µ* = 0.001.

Whereas an analytical treatment for general *L, k* and intermediate recombination rates appears to be out of reach, accurate approximations are available in the limiting case of strong recombination or of no recombination, assuming mutation rate is small. The full derivations for both cases can be found in S1 Appendix. In the following we summarize the main results.

### Strong recombination

In the limit of strong recombination we demand linkage equilibrium after each recombination step. This is satisfied if we use the so-called communal recombination scheme [58]. In this scheme an individual is not the offspring of a pair of parents. Rather, its genotype is aggregated by choosing the allele at each locus from a randomly selected parent. Hence the probability of occurrence of an allele in the offspring genotype after recombination is given by the allele frequency of the whole population, which is precisely the definition of linkage equilibrium. In order to obtain an approximation for the mutational robustness we further assume that the mutation rate *µ* is small, which in turn implies a low frequency of mutant alleles.

Following the derivation in S1 Appendix this leads us to the expression

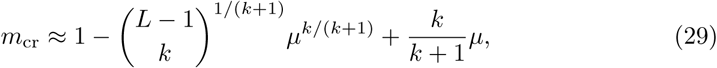

which can be approximated as

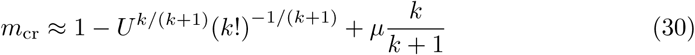

for *L* ≫ *k*, where *U* = *Lµ* is the genome-wide mutation rate and the subscript signifies the communal recombination scheme. Using Eq (29) and setting *L* = 2 and *k* = 1 we retrieve the result (20) for the two-locus model. Furthermore comparing Eq (29) and Eq (30) to numerical simulations of communal recombination illustrates their validity for large *L* (S2 Fig). If we use uniform crossover and one-point crossover instead of communal recombination, the numerical simulations suggest that the leading behaviour of 1 *− m* is still a function of *U* = *Lµ* with the same exponent *k/*(*k* + 1), which supports the universality of our findings with respect to the recombination scheme; see Fig 7.

**Fig 7.**
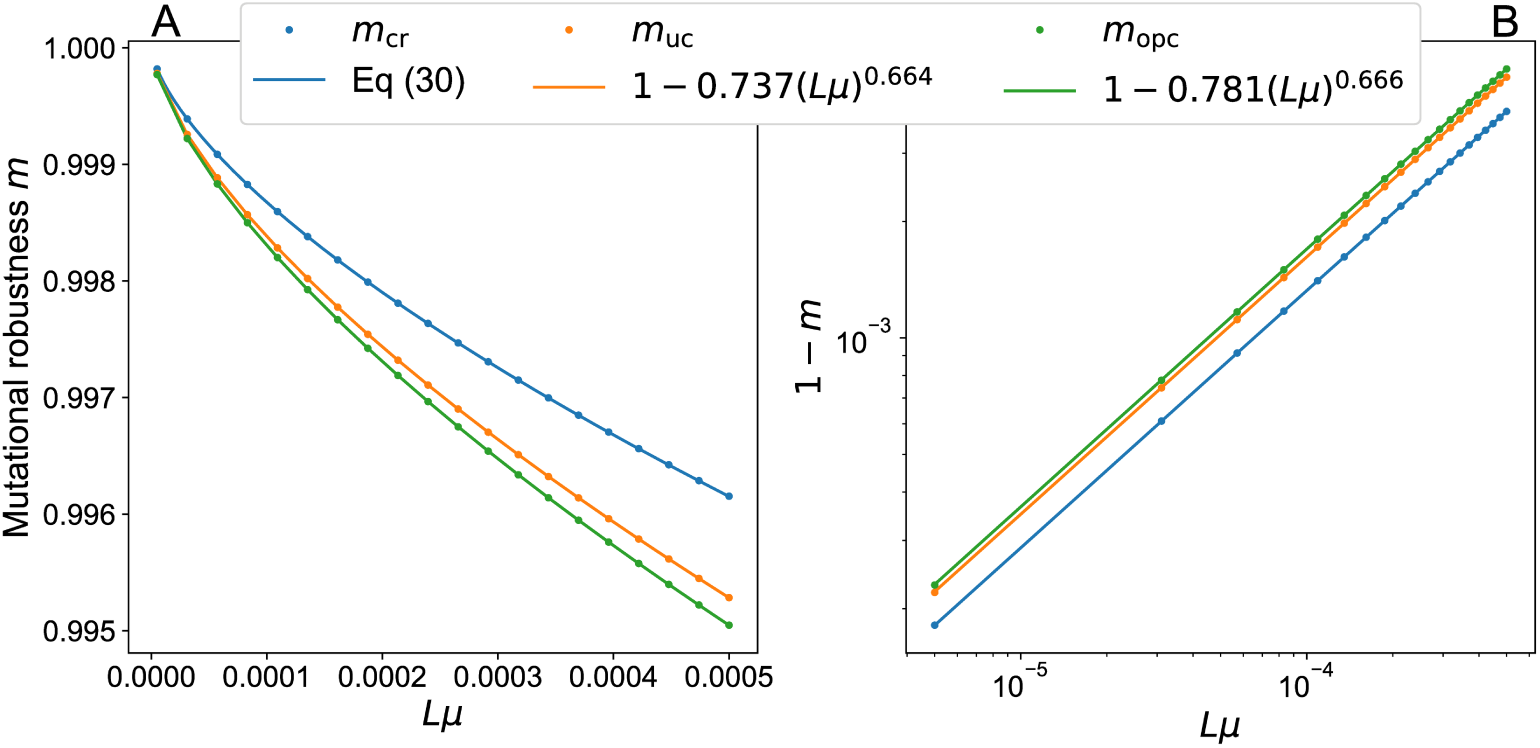
Mutational robustness in a mesa landscape with different recombination schemes. The figure compares the analytic results for communal recombination (*m*_cr_) with numerical data obtained using uniform crossover (*m*_uc_) and one-point crossover (*m*_opc_) at *r* = 1. The landscape parameters are *L* = 5, *k* = 2 and robustness is plotted as a function of the genome-wide mutation rate *Lµ*. (A) Mutational robustness on linear scales. (B) Double-logarithmic plot of 1*− m* vs. *Lµ*, illustrating the power-law behavior 1*−m* ∼(*Lµ*)^*b*^ with the exponent *b* = *k/*(*k* + 1) = 2*/*3 predicted by the analysis of the communal recombination model.

### No recombination

In order to obtain analytical results in the absence of recombination we assume that the mutation rate is small enough that only a single-point mutation occurs in one generation. This condition is fulfilled if *U* = *Lµ* ≪ 1. Interestingly, we observe that in this regime the equilibrium frequencies after selection are independent of *U.* Therefore also the mutational robustness after selection, denoted by *M*_nr_, is independent of *U.* The relation between mutational robustness after selection (*M*_nr_) and after mutation (*m*_nr_) is given by

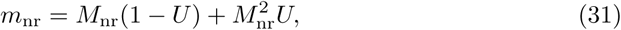

which makes it suffice to find *M*_nr_.

Assuming *k/L* ≪ 1 it is possible to link the set of stationarity conditions to the Hermite polynomials *H*_*n*_(*x*). This yields an approximation for the mutational robustness after selection as

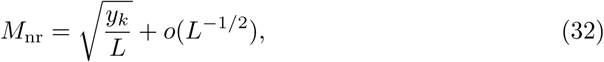

where 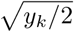 is the largest zero of *H*_*k*+1_(*x*). Correspondingly, the mutational robustness after mutation is

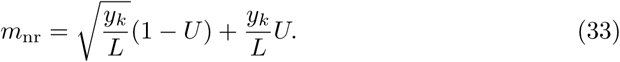

A comparison to the exact solutions for *M*_nr_, which have been obtained up to *k* = 4, confirms this approximation. If we further assume that 1 ≪ *k* ≪ *L*, we find *y*_*k*_ ∼ 4*k*, which leads to

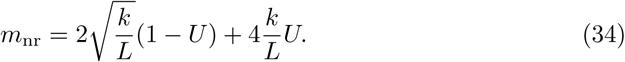

Results for the joint limit *k, L* → ∞ at fixed ratio *x* = *k/L* can be obtained from the analysis of Ref. [43], which yields

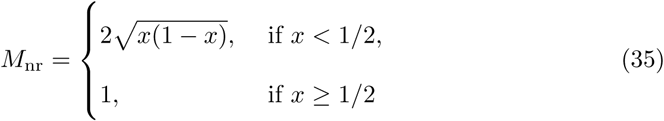

and therefore

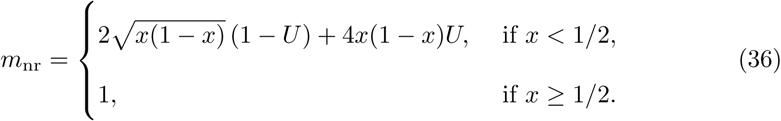

The leading behaviour for small *x* coincides with Eq (34). A comparison of these approximations to numerical solutions is given in S3 Fig.

### Comparison of the two cases

It is instructive to compare the results obtained above to the mutational robustness *m*_0_ of a uniform population distribution. For the latter we assume that all viable genotypes have the same frequency and all lethal genotypes have frequency zero. For the mesa model this yields

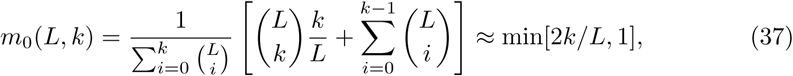

where the last approximation is valid for *L* → ∞. In Fig 8 the behavior of *m*_0_, *m*_nr_ and *m*_cr_ is depicted as a function of various model parameters. Similar to the results obtained for the two-locus model, we see that selection gives rise to a moderate increase of robustness (from 2*k/L* to 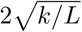 for 1 ≪ *k* ≪ *L*), but recombination has a much stronger effect and leads to values close to the maximal robustness *m* = 1 for a broad range of conditions.

**Fig 8.**
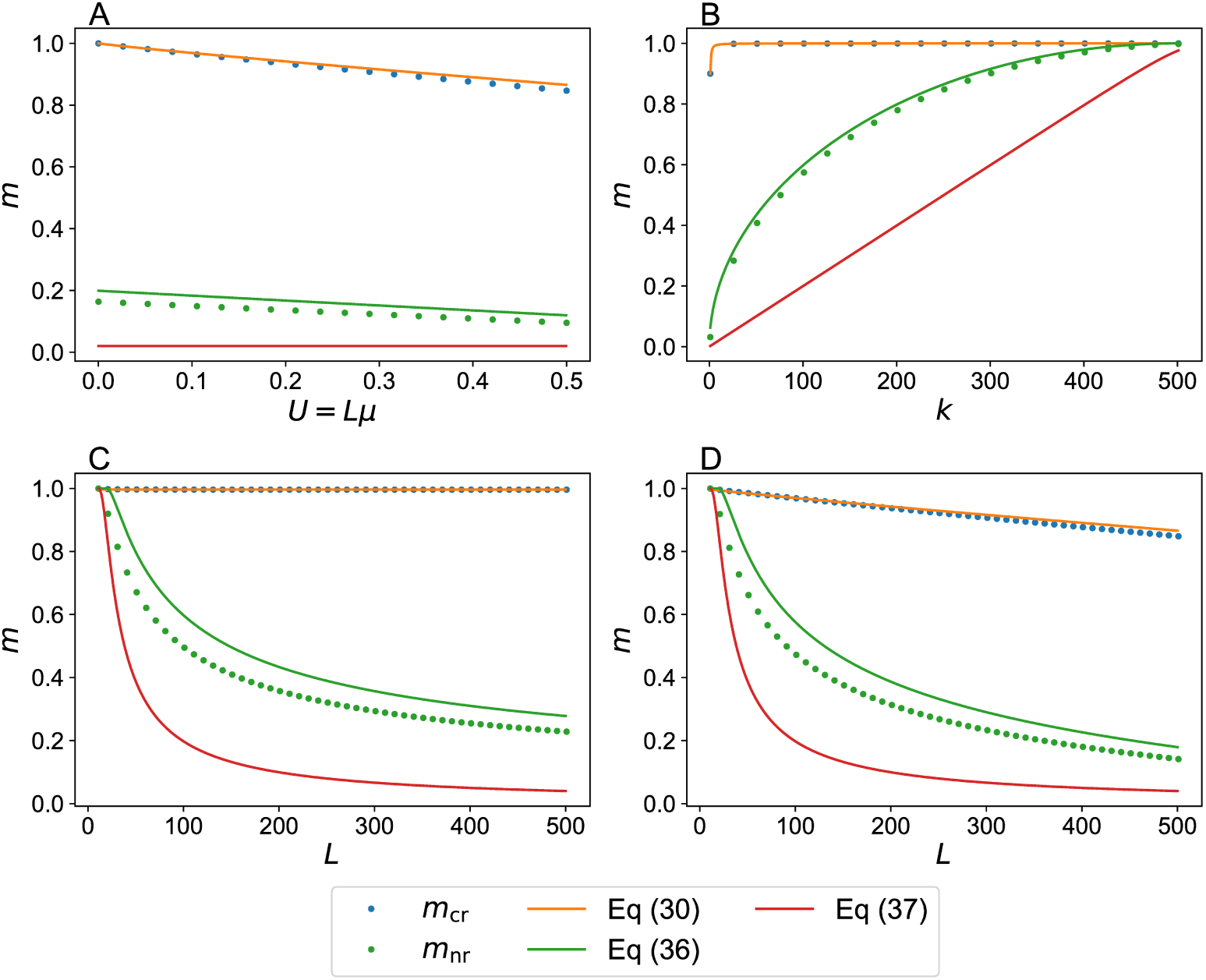
Mutational robustness in mesa landscapes with and without recombination. Numerical results for communal recombination (*m*_cr_) and no recombination (*m*_nr_) are shown as dots. The mutational robustness *m*_0_ of a uniformly distributed population, given by Eq (37), as well as the analytic expressions Eqs (30) and (36) are depicted as lines. (A) Robustness as a function of mutation rate *U* = *Lµ* for a landscape with *L* = 1000 and *k* = 10. (B) Robustness as a function of mesa width *k* at fixed *L* = 1000 and *U* = *Lµ* = 0.01. (C) Robustness as a function of genome length *L* at fixed *k* = 10 and *U* = 0.01. (D) Robustness as a function of genome length *L* at fixed *k* = 10 and *µ* = 0.001.

To elucidate the underlying mechanism, it is helpful to consider the shape of the equilibrium frequency distributions in genotype space (Fig 9). The combinatorial increase of the number of genotypes with increasing *d*_*σ*_ generates a strong entropic force that selection alone cannot efficiently counteract. As a consequence, the non-recombining population distribution is localized near the brink of the mesa at *d*_*σ*_ = *k* [43]. In contrast, the contracting property of recombination [39] allows it to localize the population in the interior of the fitness plateau where most genotypes are surrounded by viable mutants.

**Fig 9.**
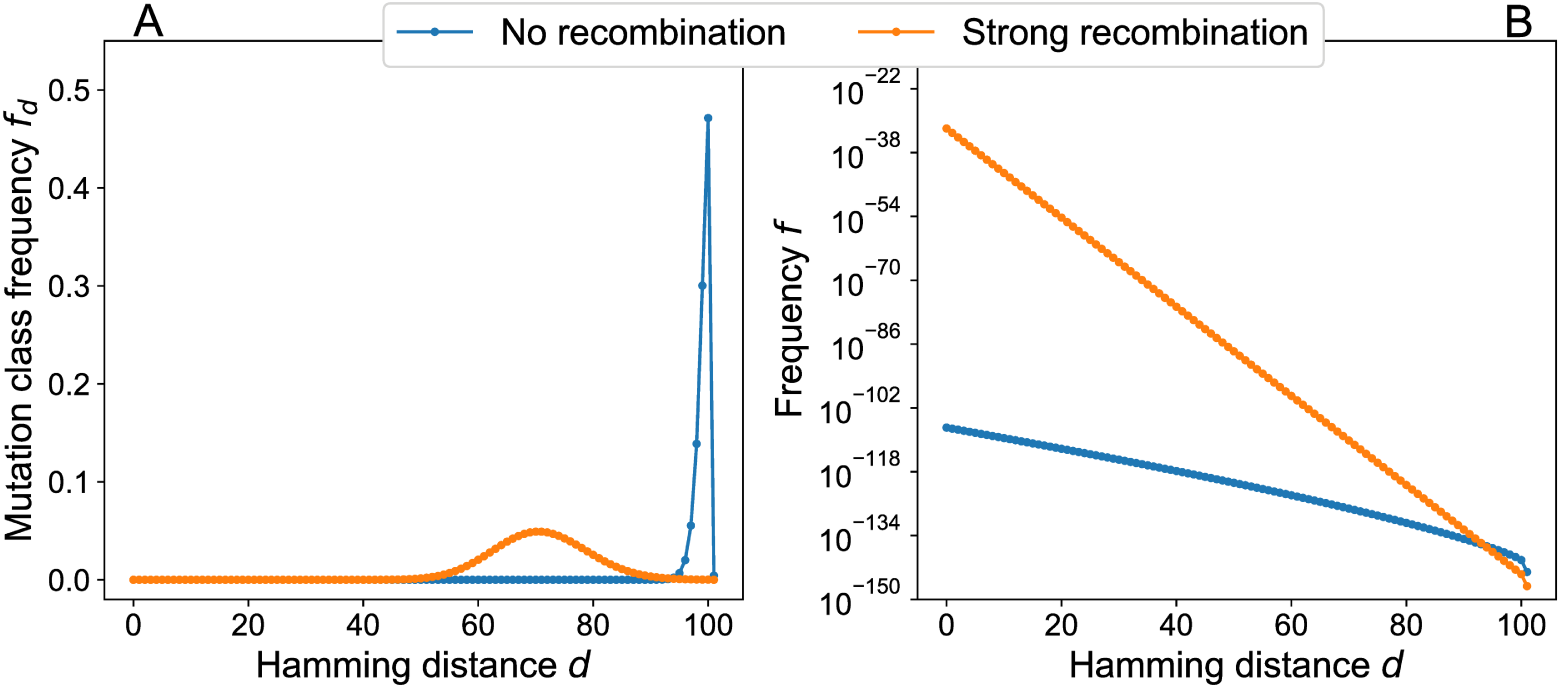
Equilibrium genotype distributions in a mesa landscape for strongly and non-recombining populations. Stationary states for populations with communal recombination and no recombination have been computed by assuming that only single point mutations occur with *U* = 0.01. Landscape parameters are *L* = 1000 and *k* = 100. The resulting mutational robustness is *m*_nr_ ≈ 0.572 for the non-recombining population and *m*_cr_ ≈1.000 for communal recombination. (A) Lumped mutation class frequencies on linear scales. In the absence of recombination the majority of the population is located at the critical Hamming distance *d* = *k*, whereas in the case of strong recombination the distribution is broader and shifed away from the brink of the mesa. (B) Genotype frequencies on semi-logarithmic scales. In both cases the genotype frequencies decrease exponentially with the Hamming distance to the wild type, but the distribution has much more weight at small distances in the case of recombination.

Fig 10 shows the corresponding recombination weight profile. Similar to the genotype frequencies in Fig 9(B) the recombination weight decays rapidly with increasing Hamming distance for r > 0, but the decay appears to be faster than exponential. Interestingly, at *d* = *k* the recombination weight decreases with increasing *r* [see also Eq (21)]. The method used to compute *λ*_*σ*_ for large mesa landscapes is explained in S1 Appendix.

**Fig 10.**
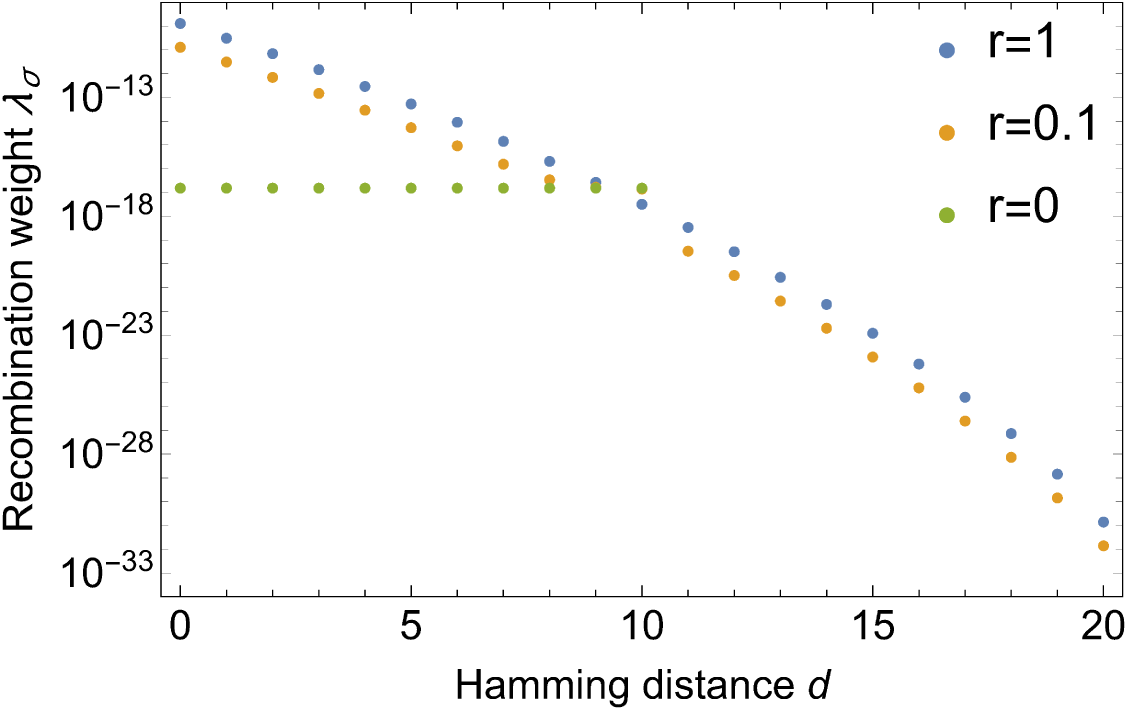
Recombination weight in a mesa landscape. The parameters of the mesa landscape are *L* = 100 and *k* = 10. For *r* = 0 the recombination weight is directly proportional to the fitness and hence equal for all viable genotypes. Already small rates of recombination are sufficient to redistribute the recombination weight such that the weight of genotypes with small Hamming distance is strongly enhanced. Beyond *d* = 20 the recombination weight is identically zero, since the recombinant of two viable genotypes cannot carry more than 2*k* mutations.

### Percolation landscapes

In the percolation landscape genotypes are randomly chosen to be viable (*w*_*σ*_ = 1) with probability *p* and lethal (*w*_*σ*_ = 0) with probability 1 *- p*. An interesting property of the percolation model is the emergence of two different landscape regimes [46, 59–61]. Above the percolation threshold *p*_*c*_, viable genotypes connected by single mutational steps form a cluster that extends over the whole landscape, whereas below *p*_*c*_ only isolated small clusters appear. Since the percolation threshold depends inversely on the sequence length,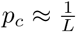, for large *L* a small fraction of viable genotypes suffices to create large neutral networks. This allows a population to evolve to distant genotypes without going through lethal regions, and correspondingly the percolation model is often used to study speciation [32, 46]. A network representation of the percolation model is shown in Fig 11. The algorithm used to generate this visual representation is explained in Models and Methods.

**Fig 11.**
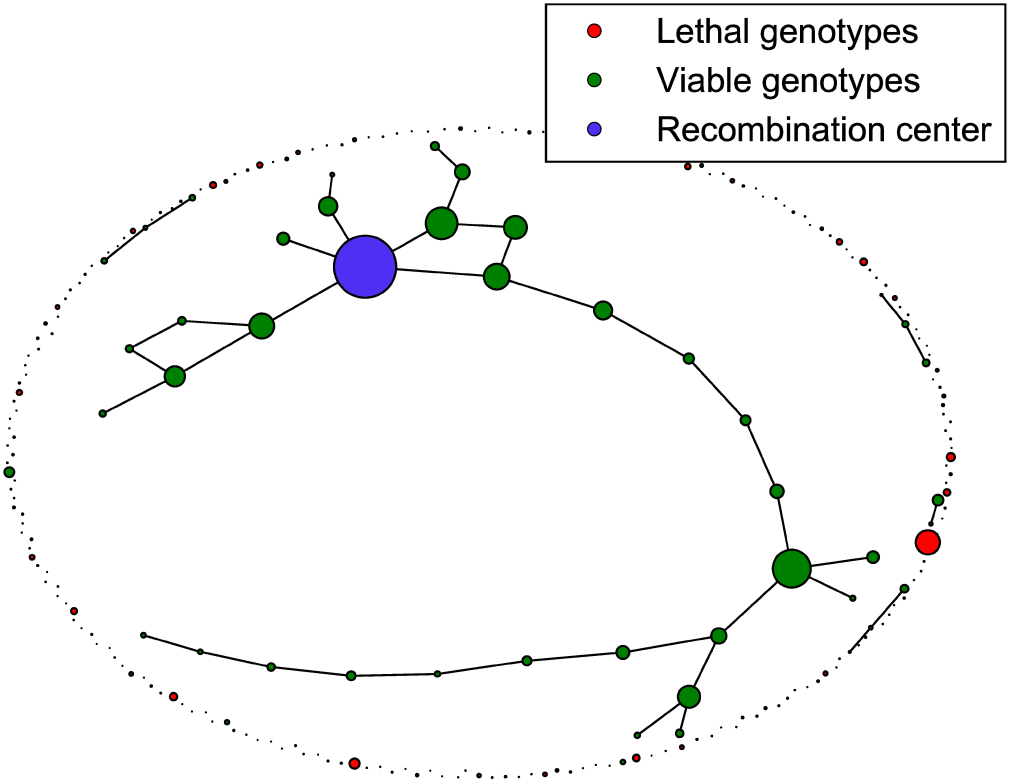
Network representation of a percolation landscape. The figure shows a percolation landscape with *L* = 8 loci and a fraction *p* = 0.2 of viable genotypes. Viable genotypes at Hamming distance *d* = 1 are connected by edges, and the node area of a genotype *σ* is proportional to 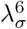, where the recombination weight *λ*_*σ*_ is defined in Eq (10). The recombination center is the genotype with the largest recombination weight.

Fig 12 shows three exemplary stationary genotype frequency distributions on the landscape depicted in Fig 11. In the absence of recombination the equilibrium frequency distribution is unique, but in the presence of recombination the non-linearity of the dynamics implies that multiple stationary states may exist [44, 51, 52]. Fig 12 displays two stationary distributions for *r* = 1 which are accessed from different initial conditions. It is visually apparent that the recombining populations are concentrated on a small number of highly connected genotypes, leading to a significant increase of mutational robustness.

**Fig 12.**
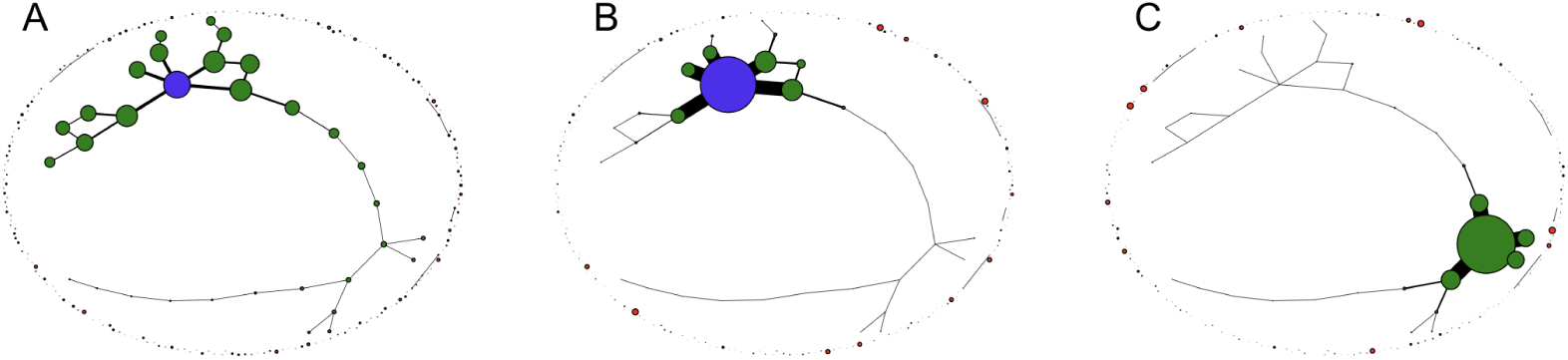
Stationary states in a percolation landscape. The figure shows three different stationary population distributions in the percolation landscape depicted in Fig 11. Node areas are proportional to the stationary frequency of the respective genotype in the population, and the edge width *e*_*σ,τ*_ between neighboring genotypes is proportional to the frequency of the more populated one, 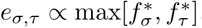. (A) Unique stationary state of a non-recombining population. (B,C) Stationary states for recombining populations undergoing uniform crossover with *r* = 1. The recombination center (purple) is the most populated genotype in (A,B), but not in (C).

To quantify this effect, the average mutational robustness 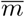 is calculated as a function of the recombination rate according to the following numerical protocol:

- A percolation landscape for given *L* and *p* is generated and the initial population is distributed uniformly among all genotypes.
- The population is evolved in the absence of recombination (*r* = 0) until the unique equilibrium frequency distribution is reached, for which the mutational robustness *m* is calculated.
- Next the recombination rate is increased by predefined increments. After increasing *r*, the population is again evolved from the stationary state obtained before increment of *r* to be the initial condition until it reaches a stationary state for which the mutational robustness is measured.
- When the recombination rate has reached *r* = 1, a new percolation landscape is generated and the process starts all over again. This is done for an adjustable number of runs over which the average is taken.

The results of such a computation are shown in Fig 13. Similar to the mesa landscapes, a strong increase of mutational robustness is observed already for small rates of recombination, and the effect is largely independent of the recombination scheme. However, in contrast to the mesa landscape the robustness does not reach its maximal value *m* = 1 for *r* = 1 and small *µ*. This reflects the fact that maximally connected genotypes with *m*_*σ*_ = 1 are very rare at this particular value of *p*.

**Fig 13.**
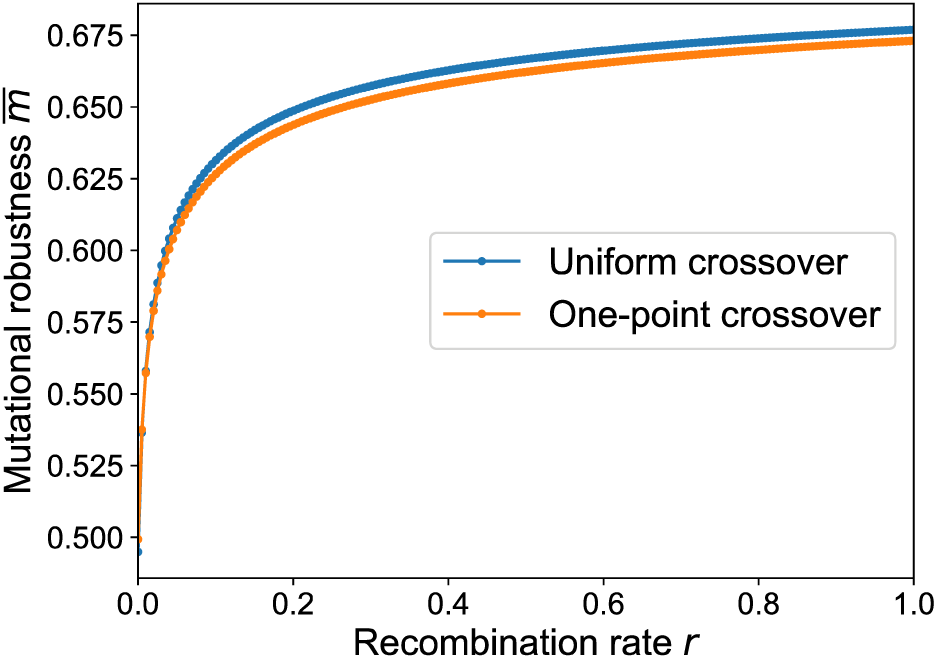
Average mutational robustness in the percolation landscape as a function of recombination rate. Mutational robustness is computed for 250 randomly generated percolation landscapes with *L* = 6 and *p* = 0.4, and the results are averaged to obtain 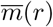. The mutation rate is *µ* = 0.001.

For the purpose of comparison we also determined the average mutational robustness 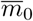 of a uniform population distribution for the percolation model. Conditioned on the number *v* of viable genotypes and assuming that *v* ≥ 1, we have *m*_0_(*v, L*) = *n*(*v, L*)*/L*, where *n*(*v, L*) is the average number of viable neighbors of a viable genotype. The latter is given by the expression

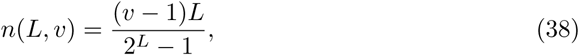

since for a given viable genotype there are *v -* 1 remaining genotypes, each of which has the probability *L/*(2^*L*^ *-* 1) to be a neighboring one. Taking into account that the number of viable genotypes is binomially distributed with parameter *p* and that the empty hypercube (*v* = 0) should yield *m*_0_ = 0 we obtain

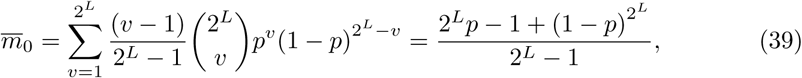

which simplifies to 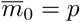 when 2^*L*^*p* ≫ 1. Note that the condition 2^*L*^*p* ≫ 1 is naturally satisfied beyond the percolation threshold.

Fig 14 illustrates that the dynamics induced by mutation and selection already increase mutational robustness compared to 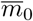 and that the addition of recombination even further increases mutational robustness for all values of *p*. The figure also displays the expected maximum number of viable neighbors of any genotype in the landscape, 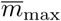, which provides an upper bound on the robustness. The fact that the numerically determined robustness remains below this bound for all *p* shows that the ability of recombination to locate the most connected genotype is limited. In S1 Appendix it is shown that 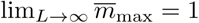 for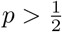.

**Fig 14.**
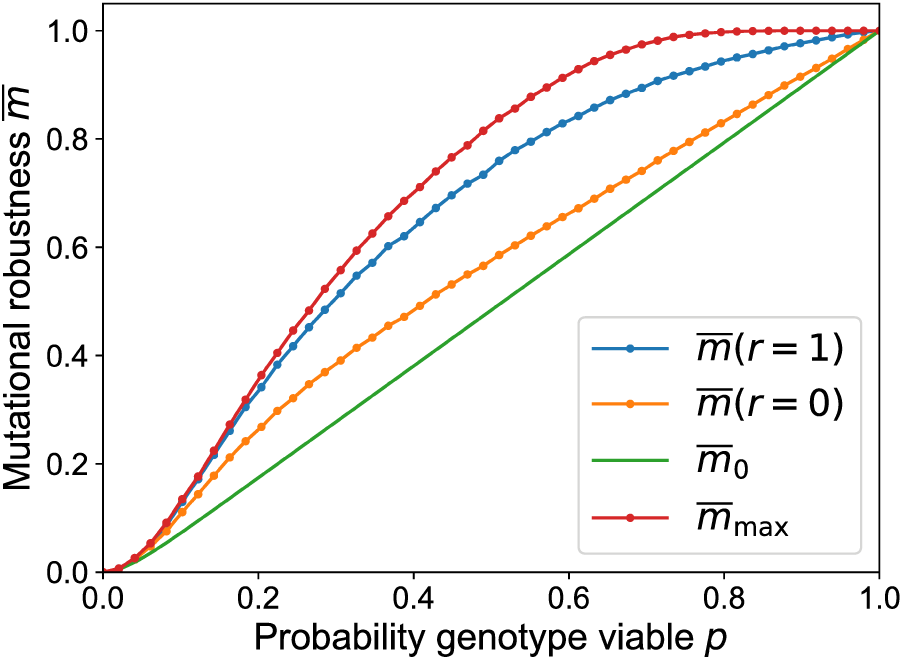
Mutational robustness in the percolation landscape as a function of the fraction of viable genotypes. The robustness for recombining 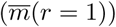 and non-recombining 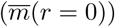 populations is obtained by averaging over 6800 randomly generated landscapes with *L* = 6 and *µ* = 0.001. In the same way the average maximal robustness 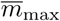 is estimated. The full line shows the analytic expression (39) for the robustness of a uniformly distributed population.

### Sea-cliff landscapes

In this section we introduce a novel class of fitness-landscape models (to be called sea-cliff landscapes) that interpolates between the mesa and percolation landscapes. Similar to the mesa landscape, the fitness values of the sea-cliff model are determined by the distance to a reference genotype *κ*^*^. The model differs from the mesa landscape in that it is not assumed that all genotypes have zero fitness beyond a certain number of mutations. Instead, the likelihood for a mutation to be lethal (to “fall off the cliff”) is taken to increase with the Hamming distance from the reference genotype. This is mathematically realized by a Heaviside step function *θ*(*x*) that contains an uncorrelated random contribution *η*_*σ*_ and the distance measure *d*(*σ, κ*^*^),

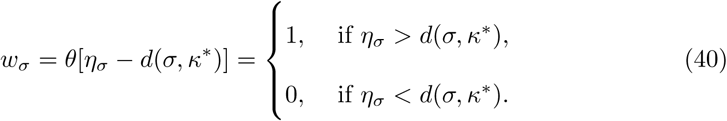

This construction is similar in spirit to the definition of the Rough-Mount-Fuji model [62, 63].

The average shape of the landscape can be tuned by the mean *c* and the standard deviation *s* of the distribution of the random variables *η*_*σ*_, which we assume to be Gaussian in the following. The average fitness at distance *d* from the reference sequence is then given by

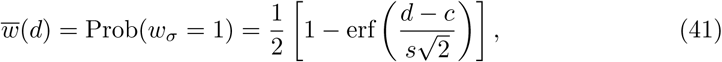

where erf(*x*) is the error function. Note that the mesa landscape is reproduced if we take *s* → ∞ limit for fixed *c* in the range *k < c < k* + 1 and the percolation landscape is reproduced if we take a joint limit *s,* |*c*| → ∞ with *c/s* fixed.

To fix *c* and *s* we introduce two distances *d*_*<*_ and and *d*_*>*_ such that 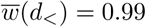 and 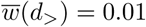, which leads to the relations

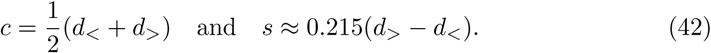

The model can be generalized to include several predefined reference sequences,

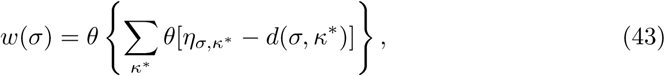

which allows to create a genotype space with several highly connected clusters. Depending on the Hamming distance between the reference sequences and the variables *c* and *s*, clusters can be isolated or connected by viable mutations.

Fig 15 shows stationary states in the absence and presence of recombination for two different sea-cliff landscapes with one and two reference genotypes, respectively. Similar to the other landscape models, mutational robustness increases strongly with recombination, due to a population concentration within a neutral cluster. In the presence of two reference genotypes the recombining population should be concentrated within a single cluster. Otherwise lethal genotypes would be predominantly created through recombination of genotypes on different clusters. This observation can also be interpreted in the context of speciation due to genetic incompatibilities [44, 46]. Without recombination genotypes on both clusters have a nonvanishing frequency, but still the larger cluster is more populated. In contrast to the percolation landscape, robustness reaches a value close to unity for large *r*, because highly connected genotypes are abundant close to the reference sequence (Fig 16).

**Fig 15.**
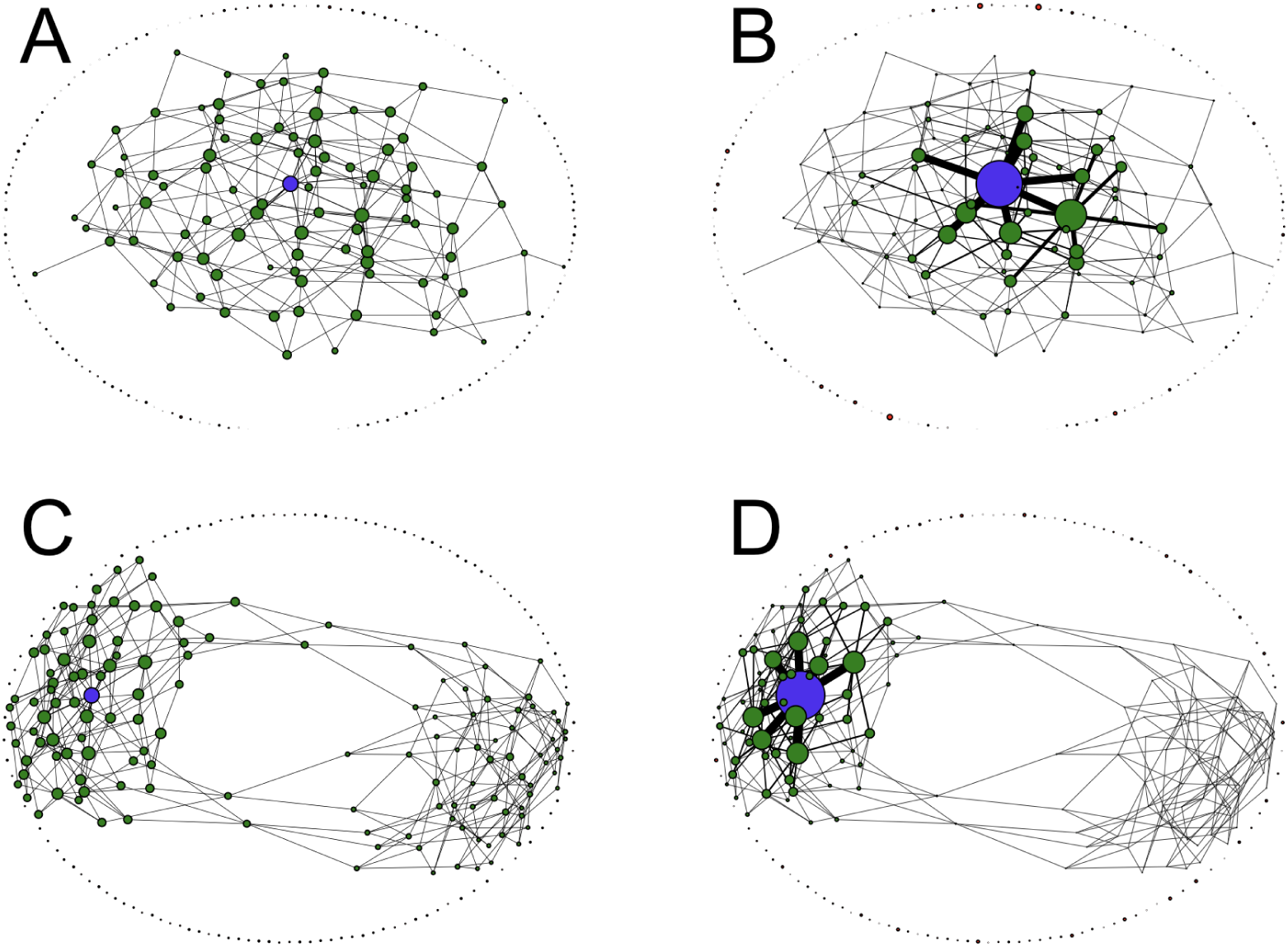
Stationary states in two different sea-cliff landscapes with and without recombination. (A,B) A single reference genotype with landscape parameters *L* = 8, *d*_*<*_ = 1 and *d*_*>*_ = 6. (C,D) Two reference genotypes which are antipodal to each other with landscape parameters *L* = 8, *d*_*<*_ = 2 and *d*_*>*_ = 4.2. (A,C) Stationary frequency distribution in the absence of recombination. (B,D) Stationary frequency distribution with uniform crossover and *r* = 1. In all cases node areas are proportional to genotype frequencies, and the recombination center is marked in blue. The edge width between neighboring genotypes is proportional to the frequency of the more populated one. The mutation rate is *µ* = 0.01.

**Fig 16.**
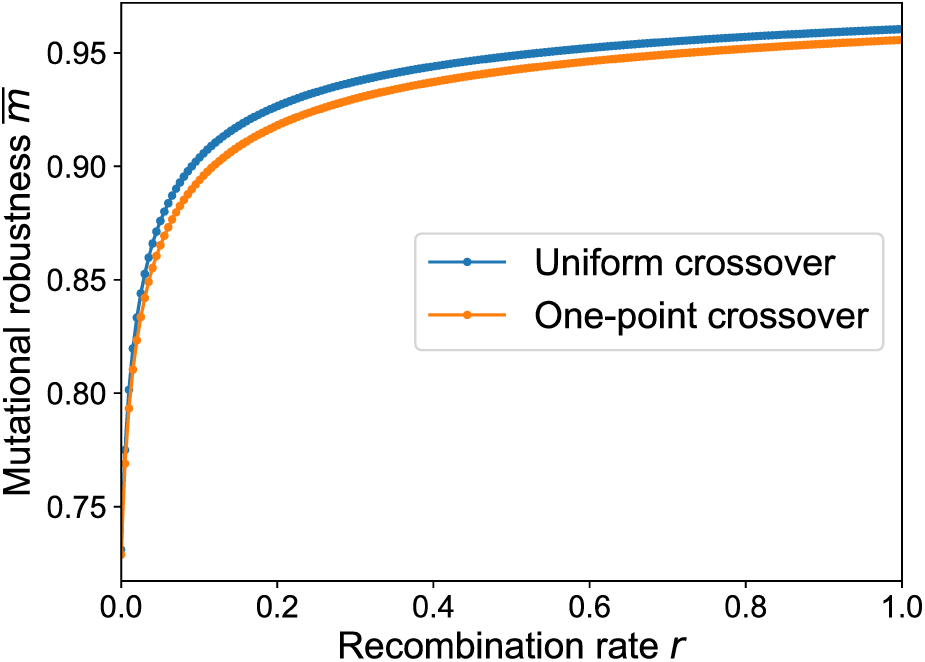
Average mutational robustness in the sea-cliff landscape as a function of recombination rate. Mutational robustness is computed for 200 randomly generated sea-cliff landscapes with parameters *L* = 6, *d*_*<*_ = 1 and *d*_*>*_ = 5, and the results are averaged to obtain 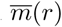. The mutation rate is *µ* = 0.001.

### Mutational robustness and recombination weight

Comparing Figs 6, 13 and 16, the dependence of mutational robustness on the recombination rate is seen to be strikingly similar. Despite the very different landscape topographies, in all cases a small amount of recombination gives rise to a massive increase in robustness compared to the non-recombining baseline. For the mesa landscape this effect can be plausibly attributed to the focusing property of recombination, which counteracts the entropic spreading towards the fitness brink and localizes the population inside the plateau of viable genotypes. In the case of the holey landscapes, however, it is not evident that focusing the population towards the center of its genotypic range will on average increase robustness, since viable and lethal genotypes are randomly interspersed.

To establish the relation between recombination and mutational robustness on the level of individual genotypes, in Fig 17 we plot the recombination weight of each genotype against its robustness *m*_*σ*_. A clear positive correlation between the two quantities is observed both for percolation and sea-cliff landscapes. Additionally we differentiate between viable and lethal genotypes. In the percolation landscape viable genotypes are uniformly distributed in the genotype space, which implies that lethal and viable genotypes have on average the same number of viable point-mutations. Nevertheless the recombination weight of viable genotypes is larger. The fitness of a genotype influences its own recombination weight, because the genotype itself is a possible outcome of a recombination event.

**Fig 17.**
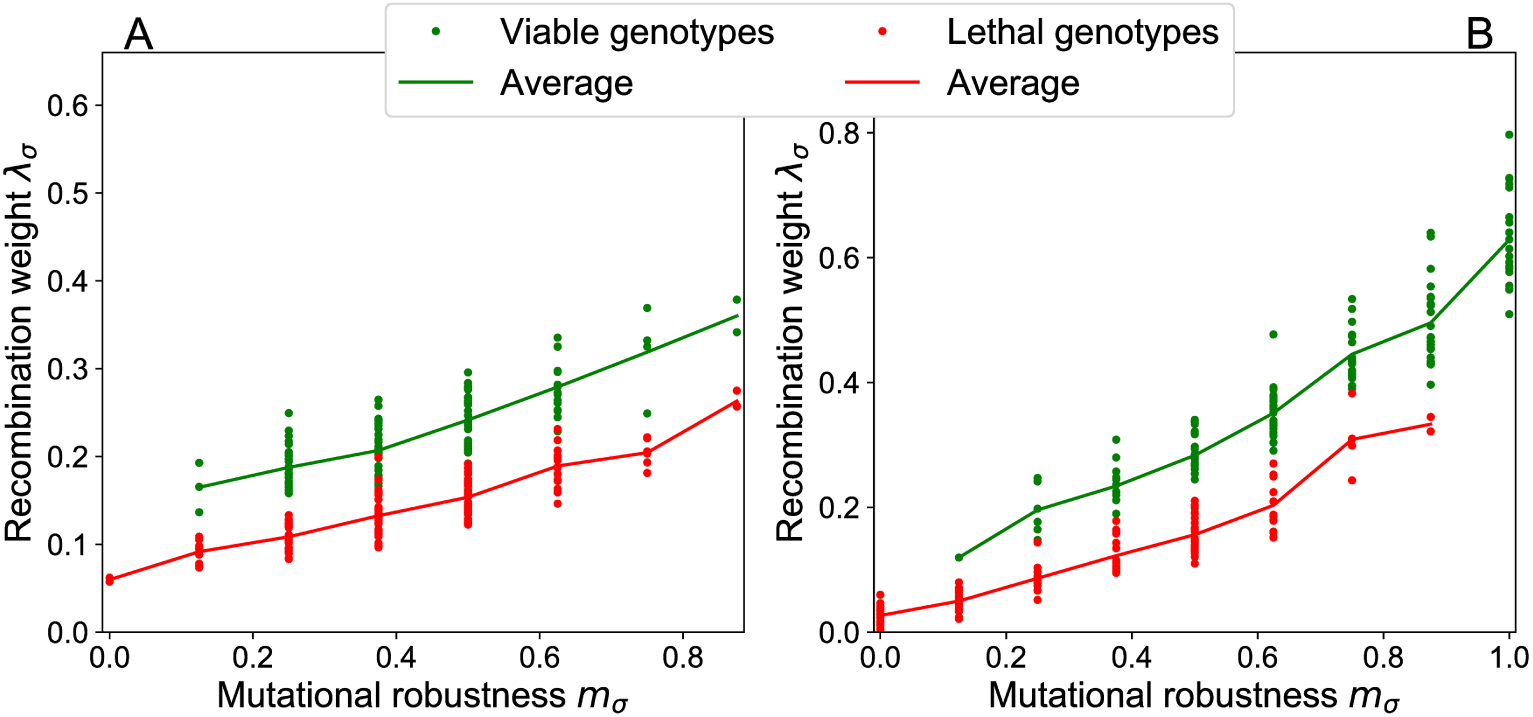
Mutational robustness correlates with recombination weight. The recombination weight of genotypes is plotted against their mutational robustness for (A) a percolation landscape with parameters *L* = 8, *p* = 0.4 and (B) a sea-cliff landscape with parameters *L* = 8, *d*_*<*_ = 2, *d*_*>*_ = 6. For the evaluation of the recombination weight (10), uniform crossover at rate *r* = 1 is assumed.

In non-neutral fitness landscapes the redistribution of the population through recombination competes with selection responding to fitness differences, and the generalized definition (11) of the recombination weight captures this interplay. To exemplify the relation between recombination weight and mutational robustness in this broader context, we use an empirical fitness landscape for the filamentary fungus *Aspergillus niger* originally obtained in [64]. In a nutshell, two strains of *A. niger* (N411 and N890) were fused to a diploid which is unstable and creates two haploids by random chromosome arrangement. Both strains are isogenic to each other, except that N890 has 8 marker mutations on different chromosomes, which were induced by low UV-radiation. Through this process 2^8^ = 256 haploid segregants can theoretically be created of which 186 were isolated in the experiment. As a result of a statistical analysis it was concluded that the missing 70 haploids have zero fitness [65].

In order to illustrate the fitness landscape, a network representation is employed where genotypes are arranged in a plane according to their fitness and their Hamming distance to the wild type, which in this case is the genotype of maximal fitness. In Figs 18A,B node sizes are adjusted to the recombination weights and mutational robustness of genotypes, respectively, in order to display the distribution of these quantities. In accordance with the analyses for neutral fitness landscapes, a clear correlation between the recombination weights and mutational robustness is shown in Fig 18C. Since fitness values are not binary we further consider the correlation between the recombination weights and fitness values (Fig 18D). The recombination center is one of the maximally robust genotypes *m*_*σ*_ = 1, but it is not the fittest within this group. The wild type has maximal fitness but, by comparison, lower robustness (*m*_*σ*_ = 7*/*8).

**Fig 18.**
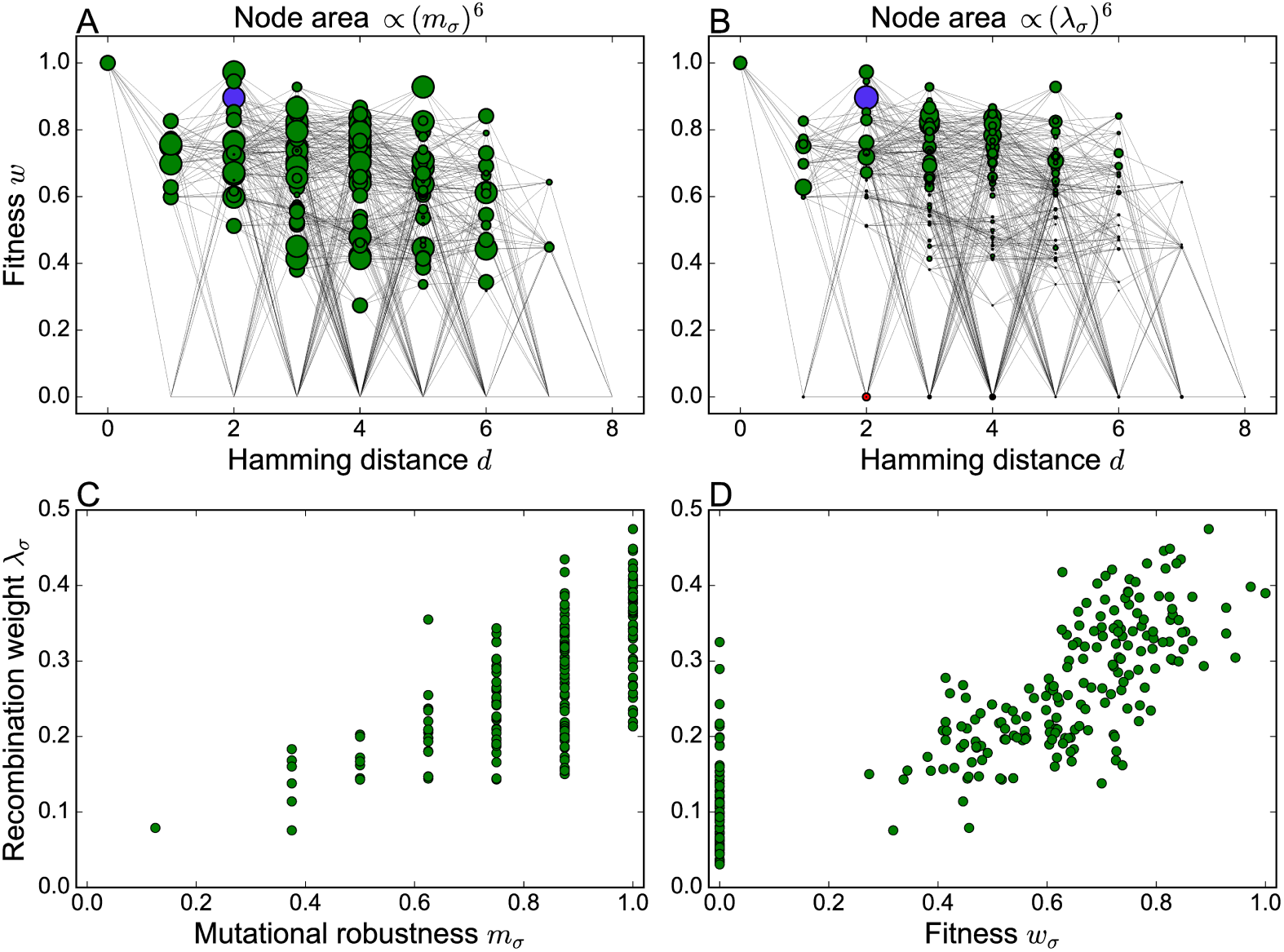
The empirical *A. niger* fitness landscape. (A,B) Two-dimensional network representation of the fitness landscape with node sizes determined by the mutational robustness *m*_*σ*_ and the recombination weight *λ*_*σ*_, respectively. In order to make the differences between genotypes more conspicuous, the node area is chosen proportional to the sixth power of these quantities. The recombination weight is evaluated for uniform crossover with *r* = 1, and the recombination center is highlighted in purple. (C,D) Recombination weight plotted against mutational robustness and genotype fitness, respectively. Lethal genotypes with *w*_*σ*_ = 0 appear only in panel D.

Fig 19 highlights how the recombination weights change as a function of the recombination rate and how this affects the stationary state of a population. For small recombination rates the recombination weight of each genotype mainly depends on its own fitness, and therefore the wild type coincides with the recombination center. With increasing recombination rate the surrounding fitness landscape topology becomes more important and the recombination center switches to a genotype at Hamming distance *d* = 2. In contrast to the numerical protocol described previously, in the simulations used to generate Figs 19C-E the population is reset to a uniform distribution before the recombination rate is increased. Otherwise the population would continue to adapt to the wild type, which has the highest fitness and from which it cannot escape because of peak trapping [20, 24]. Starting from an initially uniform distribution the population will adapt to one of three possible final genotypes which depend on the recombination rate. For small and large recombination rates the most abundant genotype is also the recombination center (Figs 19C and E), whereas for intermediate recombination rates the population chooses another genotype that is also located at Hamming distance *d* = 2 but has higher fitness (Fig 19D). The recombination center ultimately dominates the population, not only because it is maximally connected (*m*_*σ*_ = 1), but also because the genotypes that it is connected to have high fitness. In this sense the sequence of transitions in the most abundant genotype that occur with increasing recombination rate is akin to the scenario described previously in non-recombining populations as the “survival of the flattest” [43, 66]. Along this sequence mutational robustness increases monotonically whereas the average fitness of the population actually declines (S4 Fig).

**Fig 19.**
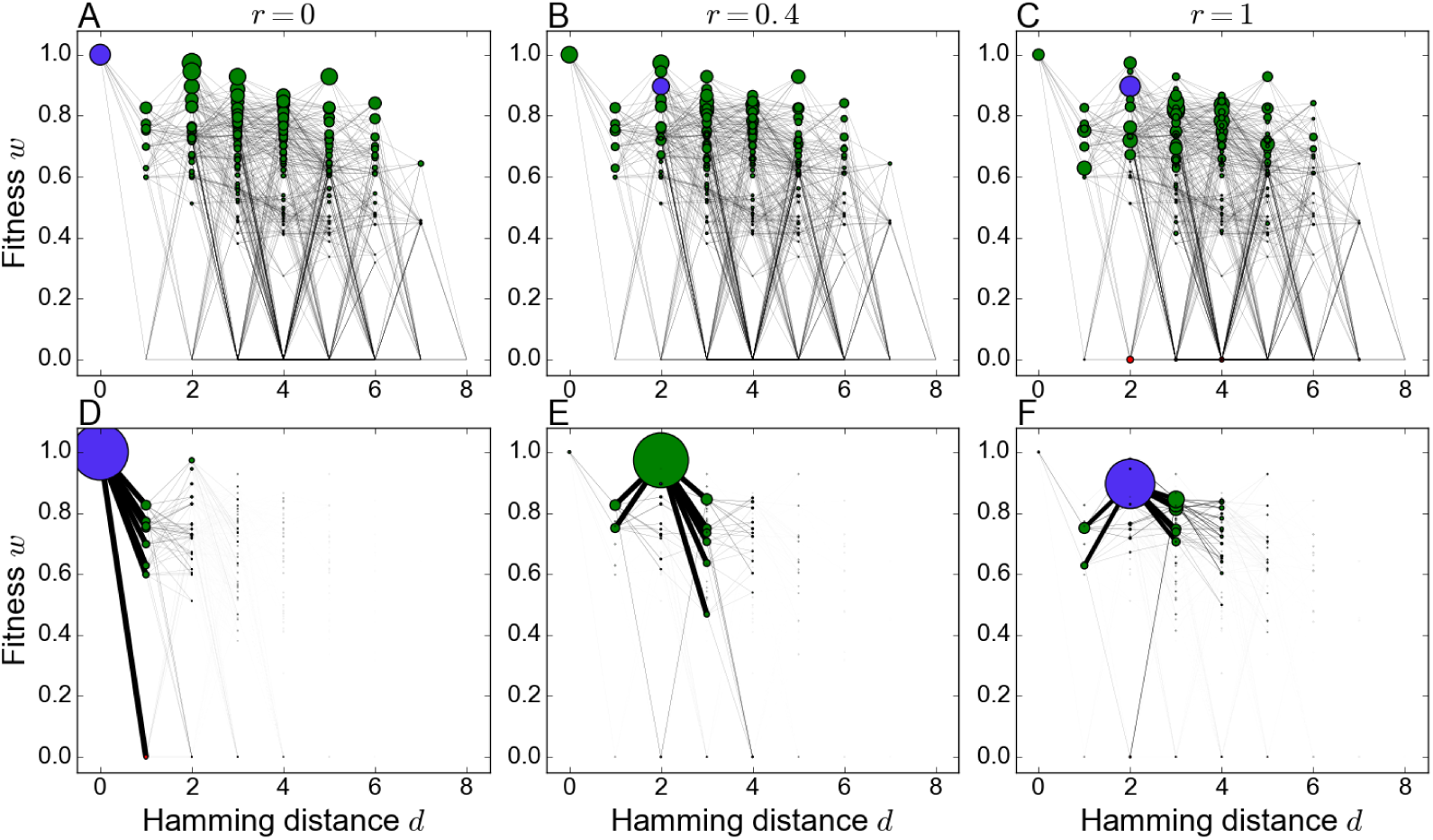
Recombination weights and stationary states at different recombination rates. (A-C) Two-dimensional network representation of the *A. niger* fitness landscape with node areas proportional to the sixth power of the recombination weight for recombination rates *r* = 0, *r* = 0.4 and *r* = 1, respectively. (D-E) Two-dimensional network representation of the *A. niger* fitness landscape with node areas proportional to the stationary genotype frequency at the same recombination rates and mutation rate *µ* = 0.005. The edge width between neighboring genotypes is proportional to the frequency of the more populated one.

## Discussion

Despite a century of research into the evolutionary bases of recombination, a general mechanism explaining the ubiquity of genetic exchange throughout the domains of life has not been found [15, 16]. Even within the idealized scenario of a population evolving in a fixed environment, whether or not recombination speeds up adaptation and leads to higher fitness levels depends in a complicated way on the structure of the fitness landscape and the parameters of the evolutionary dynamics [19–24].

The most important finding of the present work is that, by comparison, the effect of recombination on mutational robustness is much simpler and highly universal. Irrespective of the number of loci, the structure of the fitness landscape or the recombination scheme, recombination leads to a significant increase of robustness that is usually much stronger than the previously identified effect of selection [30, 31]. This suggests that the evolution of recombination may be closely linked to the evolution of robustness, and that similar selective benefits are involved in the two cases. Although the relation of robustness to evolutionary fitness is subtle and not fully understood [25], it has been convincingly argued that robustness enhances evolvability and hence becomes adaptive in changing environments [27, 29, 67]. A common perspective on recombination and robustness can help to develop novel hypotheses about the evolutionary origins of both phenomena that can be tested in future computational or empirical studies.

On a quantitative level, we have shown that robustness generally depends on the ratio of recombination to mutation rates, and that the robustness-enhancing effect saturates when *r* ≫ *µ*. This observation highlights the importance of *r/µ* as an evolutionary parameter. Interestingly, even in bacteria and archaea, which have traditionally been regarded as essentially non-recombining, the majority of species displays values of *r/µ* that are significantly larger than one [68–70]. Similarly, a recent study of the evolution of *Siphoviridae* phages revealed a ratio of recombination events to mutational substitutions of about 24 [71]. In eukaryotes this ratio is expected to be considerably higher [35]. This indicates that most organisms maintain a rate of recombination that is sufficient to reap its evolutionary benefits in terms of increased robustness.

In order to clarify the mechanism through which recombination enhances robustness, we have introduced the concept of the recombination weight, which is a measure for the likelihood of a genotype to arise from the recombination of two viable parental genotypes. The recombination weight defines a “recombination landscape” over the space of genotypes which is similar in spirit to, but distinct from, previous mathematical approaches to conceptualizing the way in which recombining populations navigate a fitness landscape [72]. It is complementary to the more commonly used notion of a recombination load, which refers to the likelihood for a viable genotype to recombine to a lethal one [36, 37]. In many cases the maximum of the recombination weight correctly predicts the most populated genotype in a recombining population at low mutation rate. Moreover, the concept generalizes to non-neutral landscapes and thus permits to address situations where selection and recombination compete.

Throughout this work the effects of genetic drift have been neglected, and it would be important to extend our analysis to finite populations. Moreover, the statistical models of neutral landscapes used here should be complemented by more realistic genotype-phenotype maps arising, for example, from the secondary structures of biopolymers such as RNA or proteins, or from simple genetic, metabolic or logical networks [27, 36, 38, 73]. Research along these lines will help to corroborate the relationship between recombination and robustness that we have sketched, and to further elucidate the origins of these two pervasive features of biological evolution.

## Acknowledgments

AK and JK acknowledge support by DFG through CRC 1310 *Predictability in evolution* and SPP 1590 *Probabilistic structures in evolution*. SCP acknowledges support by the Basic Science Research Program through the National Research Foundation of Korea (NRF) funded by the Ministry of Science and ICT (Grant No. 2017R1D1A1B03034878).

## Supporting information

**S1 Appendix.** This appendix contains detailed derivations of analytic results presented in the main text.

**S1 Fig. Population heterogeneity decreases with increasing recombination rate.** The figure shows the entropy of the genotype frequency distribution in the two-locus model defined as 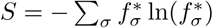. For small mutation rates the strongly recombining population primarily consists of a single genotype, which implies that *S* → 0.

**S2 Fig. Mutational robustness for the mesa landscape with communal recombination.** The figure compares the analytic approximations in Eqs (29) and (30) to the numerical solution of the stationary genotype frequency distribution for the communal recombination scheme. The two panels show the mutational robustness as a function of the genome-wide mutation rate in linear (A) and double-logarithmic (B) scales, respectively. The parameters of the mesa landscape are *L* = 30 and *k* = 3.

**S3 Fig. Mutational robustness for the mesa landscape in the absence of recombination.** The figure compares the analytic predictions in Eqs (35) and (36) to the numerical solution for the genotype frequency distribution in the absence of recombination. The two panels show the mutational robustness (A) after selection and after mutation as a function of the scaled mesa width *x*_0_ = *k/L* for *L* = 1000 and *U* = 0.01.

**S4 Fig. Mutational robustness and average fitness in the empirical *A. niger* fitness landscape.** The mutational robustness and the population-averaged fitness in the stationary state are computed as a function of recombination rate by evolving the population from a uniform initial genotype distribution at mutation rate *µ* = 0.005. Jumps mark changes in the most populated genotype.

